# Human glioblastoma cell motility depends on the activity of the cysteine metabolism enzyme 3-Mercaptopyruvate sulfurtransferase

**DOI:** 10.1101/2022.01.21.477271

**Authors:** Mirca S. Saurty-Seerunghen, Thomas Daubon, Léa Bellenger, Virgile Delaunay, Gloria Castro, Joris Guyon, Ahmed Rezk, Sylvie Fabrega, Ahmed Idbaih, Fabien Almairac, Fanny Burel-Vandenbos, Laurent Turchi, Thierry Virolle, Jean-Michel Peyrin, Christophe Antoniewski, Hervé Chneiweiss, Elias A. El-Habr, Marie-Pierre Junier

## Abstract

Cancer cells in similar functional states are found in all glioblastoma, despite the genomic heterogeneity observed between and within these brain tumors. Metabolism being downstream of all signaling pathways regulating cell behaviors, we looked for metabolic weaknesses in link with motility, a key functional state for glioblastoma aggressiveness. A signature-driven data reduction approach highlighted motile cells present in thirty tumors from four independent single-cell transcriptomic datasets. Analyses integrating trajectory modeling disclosed, as characteristic of motile cells, enhanced oxidative stress coupled with mobilization of the cysteine metabolism enzyme 3-Mercaptopyruvate sulfurtransferase (MPST). The soundness of this prediction was verified using migration and invasion assays with patient-derived cells and tissue organoids. Pharmacological and genetic manipulations showed that enhanced ROS production and MPST activity are required for the cells’ motility. Biochemical assays indicated that MPST acts by protecting protein cysteine residues from dismal hyperoxidation. In vivo, MPST knockdown translated in reduced tumor burden, and a robust increase in mice survival. These results show that enhanced oxidative stress coupled with MPST mobilization plays a key role in glioblastoma cell motility.

## Introduction

Glioblastoma (GB) cell heterogeneity is now recognized as a major issue to overcome in order to develop more efficient therapies for this primary brain tumor that remains with a very poor prognosis. The median survival of patients does not exceed twenty months despite aggressive treatments combining surgical removal of the tumor with chemotherapy and radiotherapy (Stupp et al 2017). GB is a paradigm of intra-tumor heterogeneity. Normal and cancerous cells intermingle, and tumor territories differing in tumor cell densities, vascular perfusion and necrosis, entangle (Hambardzumyan & Bergers 2015). Genome-wide DNA and RNA sequencing at the tissue and single cell levels have further illustrated GB heterogeneity. They showed the coexistence, within each tumor, of cancer cells bearing differing mutational loads and genomic rearrangements and varying ontogenic similarities with neuro-developmental lineages (Snuderl et al 2011) (Sottoriva et al 2013) ((Lee et al 2017), (Puchalski et al 2018) (Neftel et al 2019) (Couturier et al 2020) (Bhaduri et al 2020) (Wang et al 2020) (Richards et al 2021). Notwithstanding these variable mutation loads, chromosome rearrangements, and developmental-like profiles, we reasoned that GB heterogeneity would appear less overwhelming when considering the behavior or functional state of tumor cells in link with their underlying metabolic pathways. Tumor cells in identical functional states can be detected within each GB and across patients, regardless of their genomic backgrounds. Downstream of all signaling pathways regulating cell functional states is metabolism. Studies have shown that changes in metabolic activity not only accompany changes in cell functional states but can also drive changes as well (El-Habr et al 2017) (Flavahan et al 2013) (Oizel et al 2017). Thus, metabolic enzymes playing key roles in the adoption and/or maintenance of cell functional states crucial for tumor development, as observed at the time of patients’ diagnosis, constitute therapeutic targets able to overcome tumor heterogeneity.

Cell motility is a key contributor to GB malignancy and poor prognosis. GB cells are highly infiltrative, making complete surgical resection of the tumor impossible. Moreover, currently used irradiation and chemical therapies have been shown to promote GB cell invasiveness (Huber et al 2013) (Lu et al 2012). Cell motility encompasses two processes: migration, i.e. the cell’s ability “to move around in a space that is freely available”, and invasion, a process involving microenvironment remodeling by the cells (de Gooijer et al 2018). Motile cells have to develop protrusions in the direction of migration, disrupt adhesion sites at the cell rear, degrade and remodel the extracellular matrix (ECM) and ultimately make new connections at the cell front by remodeling their cytoskeleton so that they are pushed forward, making cell motility a highly integrated process (Armento et al 2017). Three major migration/invasion routes have been described: the narrow and tortuous extracellular space of the brain parenchyma (Scherer 1938), the perivascular spaces surrounding blood vessels (Scherer 1938) (Montana & Sontheimer 2011) (Watkins et al 2014) (Cuddapah et al 2014) and white matter tracts, notably the corpus callosum, used as a highway for bilateral brain hemisphere invasion (Scherer 1938) (Pedersen et al 1995). Both single cell and collective migration have been reported (Vollmann-Zwerenz et al 2020) (Volovetz et al 2020), the latter appearing to be facilitated by the formation of an interconnected network of tumor microtubes (Osswald et al 2015). So far, an integrated view of the metabolic pathways at play in a motile GB cell is lacking.

Here, we applied a signature-driven data reduction approach to highlight subpopulations of cells in a motile state present in thirty tumors from four independent single-cell transcriptomic datasets. We previously demonstrated that such an approach allows to uncover cell functional states and their associated key metabolic players from transcriptomes of single cells derived from surgical resections of the patients’ tumors (Saurty-Seerunghen et al 2019). Computational analyses integrating cell trajectory modeling characterized cells with high motile potential as endowed with higher energetic needs and oxidative stress than their more static counterparts, and highlighted the 3-Mercaptopyruvate sulfurtransferase (MPST) enzyme at the crossroad of the path from low to high motility. Requirement of this element of the cysteine metabolism for GB cell motility was experimentally demonstrated. Mechanistically, MPST acts by protecting protein cysteine residues from hyperoxidation. Our findings uncover a previously unsuspected involvement of MPST in the switch from low to high cell motility and maintenance of GB cell motility.

## Results

### Grouping cells according to their potential motile state

We used four independent publicly available single-cell transcriptome datasets generated from surgical resections of patients bearing IDH-wild-type GB. Two were obtained with the SMART-seq2 technology (Neftel et al 2019) (Darmanis et al 2017) and two with the 10X Genomics technology (Neftel et al 2019) (Pombo Antunes et al 2021). They are hereafter designated as N-S, D-S, N-10X and PA-10X, according to the initials of the first author of the paper reporting its first description and the technology used. The computational analyses were first performed using the N-S dataset, and the robustness of the results evaluated by repeating the analyses with each of the other three datasets. This choice stems from the deeper sequencing depth of SMART-seq2 data compared to 10X Genomics data, and the higher number of malignant cells and distinct patient tumors in the N-S dataset compared to the D-S dataset.

For clustering cells in a motile state, we implemented a data reduction approach driven by a molecular signature (Fig.1a), a previously reported approach grouping cells from distinct tumors according to their functional state (Saurty-Seerunghen et al 2019). Here, cells were grouped based on the combined expression levels of each gene in a motility signature, using the HCPC algorithm (Hierarchical Clustering on Principal Components) (Husson et al 2010) modified to integrate also UMAP components (Uniform Manifold Approximation and Projection) (Supplementary Figure 1A). The genes composing the motility signature were selected based on the *in vitro* and *in vivo* experimental demonstration of their requirement for GB cell motility using recent patient-derived GB cells (PDC) cultured in defined media. Thus, these inducers and effectors of GB cell motility are involved in pro-motile autocrine signaling (TGFB1, SMAD3, THBS1 (Joseph et al 2014) (Daubon et al 2019) (Joseph et al 2021)), morphological rearrangements undertaken by motile cells and linking extracellular guidance signals with the remodeling of the cytoskeleton (ACTN4 (Sen et al 2009) (Ji et al 2019) (Tentler et al 2019), PTK2 (Xia et al 2016) (Frolov et al 2016), PXN (Frolov et al 2016) (Sun et al 2017) (Lopez-Colome et al 2017), TLN1 (Kang et al 2015), VCL (Xia et al 2016)) and the remodeling of the matri-cellular environment (TNC (Xia et al 2016), SPARCL1 (Gagliardi et al 2017)). Expression of each gene of the signature is most often positively correlated with expression of the other signature genes across all cells (Fig.1b, Supplementary Table 1).

**Fig. 1.**
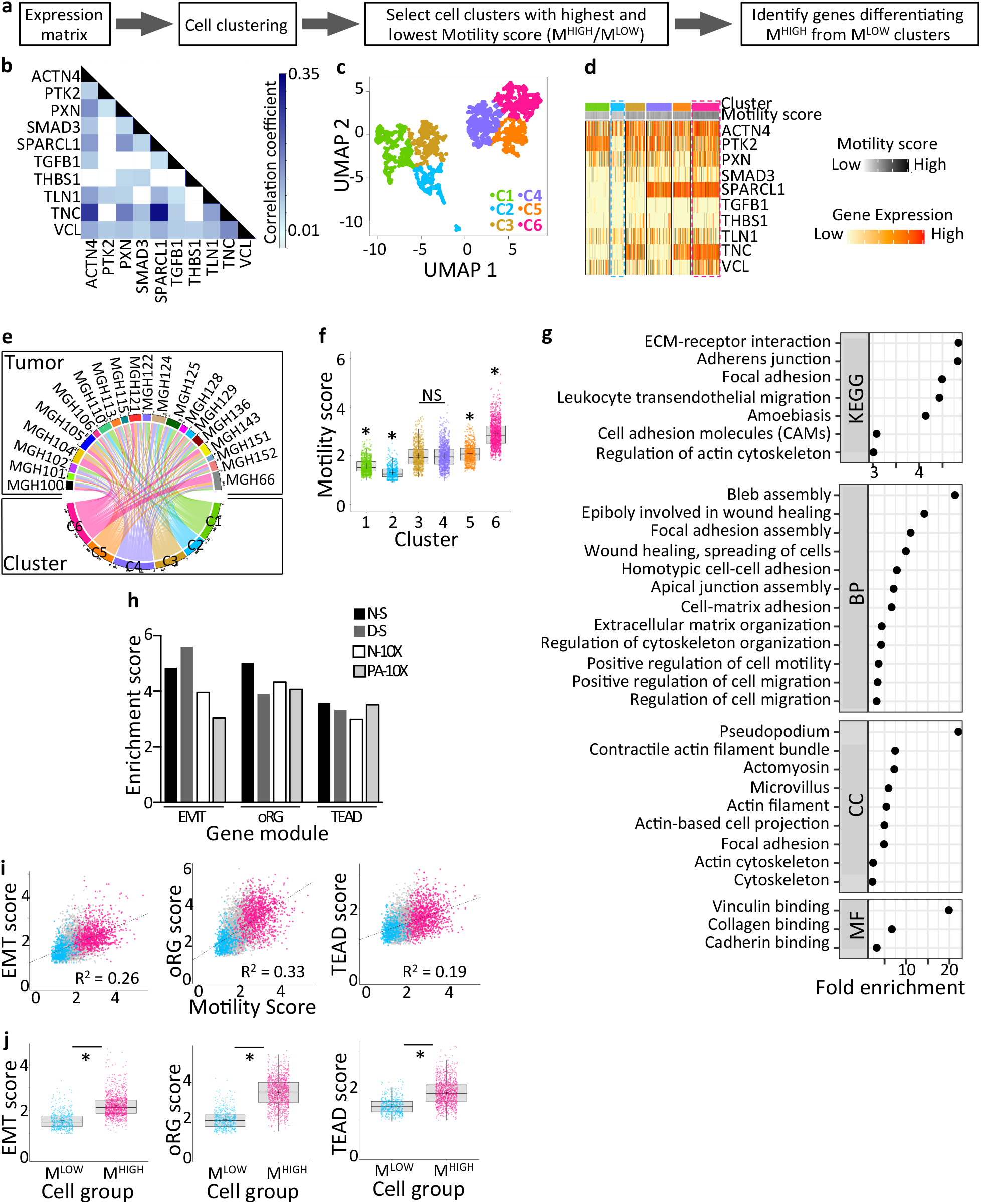
Capturing glioblastoma cells with high and low motile potential. **a**. Schematic outline of the computational analytical strategy. See text for details. **b**. Correlations between expression of the ten genes of the motility signature. Only significant correlations are shown (p < 0.01). **c**. Malignant cell clustering based on the motility signature. UMAP representation. **d**. Different combinatorial expressions of the signature genes characterize each cluster. Heatmap representation of the relative expression of each signature element per cluster. **e**. Contribution of each tumor to identified clusters. Each cell cluster contains cells coming from distinct tumors (NMI score = 0.11). **f**. Identification of cell groups with the highest and lowest mean motility scores (C6: M^HIGH^, C2: M^LOW^). *: Clusters with mean motility score statistically different from each of the other clusters, p-value < 0.01, one-way ANOVA, Tukey’s multiple comparisons test. NS: non-significant. M: Motile potential of Glioblastoma cells. **g**. Motility-related terms highlighted by ontology analysis of genes overexpressed in M^HIGH^ versus M^LOW^ cells. Dot plot representation. Genes overexpressed with fold change ≥2. Benjamini-Hochberg (BH)-adjusted p-value < 0.05. KEGG: Kyoto Encyclopedia of Genes and Genomes, BP: Biological processes, CC: Cellular components, MF: Molecular functions. **h**. Enrichment in gene modules previously associated with GB cell motility in M^HIGH^ cells. EMT: genes associated with epithelio-mesenchymal transformation and GB cell motility. oRG: genes signing for outer Radial Glia (oRG)-like malignant cell population with increased invasive behavior in GB. TEAD: TEAD1-regulated genes involved in GB cell motility. p < 0.001, hypergeometric test. **i.** Linear regression model between motility score and EMT, oRG and TEAD scores. p < 0.0001. **j.** Higher EMT, oRG and TEAD scores in M^HIGH^ versus M^LOW^ cells. *: p < 0.0001, Mann-Whitney test. **b-g**, and **i-j**: Results from N-S dataset analysis.

This clustering analysis resulted in six cell clusters (Fig.1c) with distinct combinations of expression of the signature genes (Fig.1d). Of note, hierarchical clustering of UMAP components led to groups of cells with transcriptomic profiles more homogeneous than when using Hierarchical Clustering of Principal Components (HCPC), as shown by increased Silhouette width (Supplementary Figure 1B). The Normalized Mutual Information (NMI) value was then calculated to determine the contribution of different tumors to each cluster, in order to demonstrate that cell clustering is driven by functional state rather than tumor of origin as described (Saurty-Seerunghen et al 2019). A NMI value of 1 implies that clusters gather cells corresponding to a single tumor, whereas a value of 0 denotes that each tumor contributes to each cluster. As indicated by the low NMI value (NMI = 0.11), each cluster was composed of cells coming from distinct tumors (Fig.1e). Using the geometric mean of the signature gene expression levels as a motility score, we retained the two cell clusters with the highest (M^HIGH^) and lowest (M^LOW^) motile potential, respectively (Fig.1f). Differential expression analysis between M^HIGH^ and M^LOW^ cell groups (Mann Whitney, BH-adjusted p-value < 0.01) highlighted the genes overexpressed in M^HIGH^ cells (Supplementary Table 2-sheet 2).

### Enrichment of motility-related genes in M^HIGH^ cells

We evaluated the coherence of our findings by confronting them with previous experimentally-acquired knowledge. In gene ontology (GO) enrichment analyses, we found an enrichment of motility-related terms among the genes overexpressed at least two-fold in M^HIGH^ cells (log_2_ fold-change≥2), providing a first support to the relevance of our approach (Fig.1g, Supplementary Table 3-sheet2). To further probe this relevance, we applied the same analytical strategy to the other three selected datasets. THBS1 being detected in <1% malignant cells in the PA-10X dataset, it was excluded from the motility signature for that dataset. Each of these analyses resulted in clusters composed of cells coming from different tumors (NMI =0.058-0.15) (Supplementary Figure 1C-E). Differential expression analysis between M^HIGH^ and M^LOW^ cell clusters (Mann Whitney, BH-adjusted p-value < 0.01) (Supplementary Table 2-Sheet3-5), followed by GO enrichment analyses of the most overexpressed genes, again highlighted enrichment in motility-related terms (Supplementary Figure 2, Supplementary Table 3-sheet3-5). To determine the similarities of M^HIGH^ cell transcriptional profiles from distinct datasets, we compared the lists of their overexpressed genes. The highest overlaps were found between lists of genes derived from datasets obtained with the same sequencing technology. 69.2% (4852/7010) of the genes overexpressed in M^HIGH^ cells from N-S dataset were also overexpressed in M^HIGH^ cells from D-S dataset, and 86.0% (6627/7707) of the genes overexpressed in M^HIGH^ cells from N-10X dataset were also overexpressed in M^HIGH^ cells from PA-10X dataset (Supplementary Figure 1F). Comparing the four lists of genes overexpressed in M^HIGH^ GB cells, we found that 43 to 53% of the genes overexpressed in M^HIGH^ GB cells from one dataset were also overexpressed in M^HIGH^ cells from the three other datasets (Supplementary Figure 1F, Supplementary Table 2-Sheet6). These results indicate that our analytical strategy captures cells with similar transcriptional profiles in distinct datasets.

To further determine the extent to which our analytical approach captures cells in a motile state, we selected three gene modules previously found to be associated with GB cell motility (Supplementary Table 2-Sheet7), and determined their enrichment in M^HIGH^ cells and their correlation with the motility score. Most motile cells, including GB cells have been shown to undergo transcriptional reprogramming akin to the so-called epithelial-mesenchymal transformation (EMT) process. We therefore used an EMT gene module of 14 genes associated with mesenchymal transformation of neural cells and GB cell motility (Carro et al 2010) (Armento et al 2017). We also sought for enrichments in two other gene modules mobilized in invasive GB cells. One consists of 36 genes highlighted by Bhaduri and colleagues with highly correlated expression in an outer radial glia (oRG)-like malignant cell population with increased invasive behavior in GB (Bhaduri et al 2020). The other consists of 32 genes involved in the regulation of GB cell motility and driven by TEAD transcription factors (Tome-Garcia et al 2018). Each of these three gene modules was found to be enriched in M^HIGH^ cells in all analyzed dataset (Fig.1h). We also observed a positive linear relationship between the motility score and the EMT, oRG and TEAD scores (Fig.1i, Supplementary Figure 3). As expected from these correlations, the EMT, oRG and TEAD scores were significantly higher in M^HIGH^ cells compared to M^LOW^ cells from each dataset (Fig.1j, Supplementary Figure 3).

Altogether, these results support the relevance of the analytical strategy implemented for capturing cells with high motile potential from distinct datasets.

### Metabolic characteristics of cells with high motile potential

We went on to identify metabolic pathways deregulated in GB cells with high motile potential. The genes coding for enzymatic components of the metabolic pathways overexpressed in M^HIGH^ cells were identified in each dataset using the 2019 KEGG list of 1586 metabolic elements (Kanehisa et al 2017) (Supplementary Table 4-Sheet2). KEGG pathway enrichment analyses highlighted metabolic pathways related to extracellular matrix (ECM) modeling (e.g. glycosaminoglycan synthesis/degradation) and membrane composition rearrangements (e.g. unsaturated fatty acids (FA), FA elongation, GPI-anchor biosynthesis) (Fig.2a, Supplementary Table 4-Sheet3-6). In addition to these metabolic pathways expected to be active in motile cells, we also observed enrichment in pathways involved in energy production (Fig.2a, Supplementary Table 4-Sheet3-6). The upregulation of genes involved in glycolysis, amino acid catabolism, fatty acid degradation, protein recycling (lysosome), TCA cycle, oxidative phosphorylation, suggests that GB cells with high motile potential have higher energetic needs than cells with low motile potential. Enrichment in oxidative phosphorylation, a term gathering elements of the electron transport chain (ETC), associated with enrichment in TCA cycle components, suggests in addition a higher mitochondrial load in M^HIGH^ cells. Finally, we observed enrichment in pathways involved in anti-oxidative processes, such as the pentose phosphate pathway (PPP), and the cysteine, sulfur and glutathione metabolisms (Fig.2a, Supplementary Table 4-Sheet3-6). Enrichment in pathways or organelles involved in ROS scavenging and/or production of anti-oxidants (porphyrin and selenocompound metabolisms, peroxisome), as well as enrichment in purine/pyrimidine metabolism, required for DNA/RNA repair, are also compatible with enhanced responses to oxidative stress. Overall, the results of this analysis point to an enhanced mobilization of anti-oxidative pathways counteracting the deleterious actions of oxidative compounds derived from the over-mobilization of energy production pathways.

**Fig. 2.**
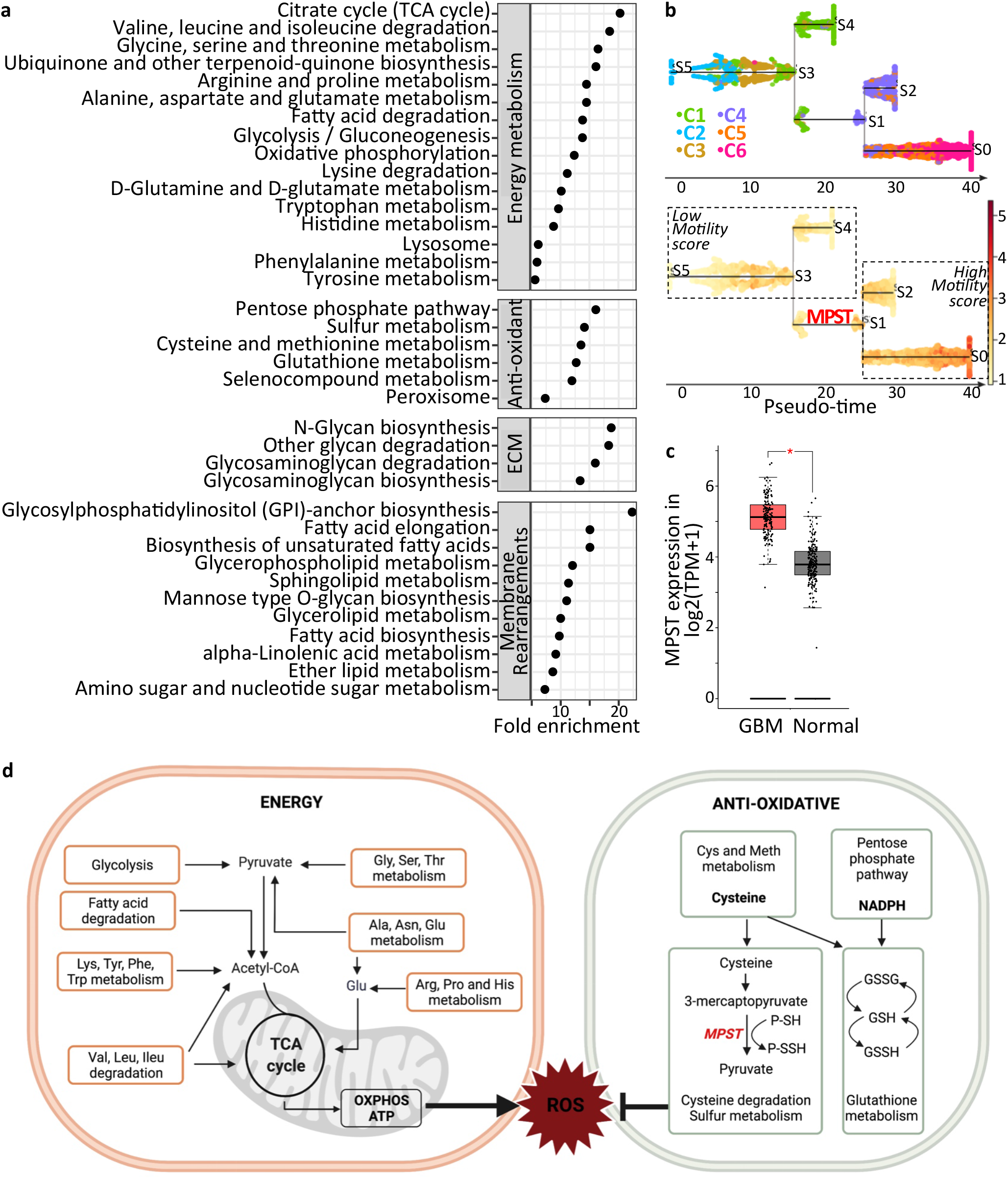
Metabolic characteristics of glioblastoma cells with high motile potential. **a.** KEGG pathway analysis of metabolism genes overexpressed in M^HIGH^ cells compared to M^LOW^ cells highlights enrichment in metabolic pathways involved in energy production, oxidative stress response, extracellular matrix (ECM) modeling and membrane composition rearrangements. BH-adjusted p-value < 0.05. Results from N-S dataset analysis. **b.** Inferred trajectory of glioblastoma cells from low to high motility. Cells colored by motility clusters (top panel) and motility score (bottom panel). MPST (Mercaptopyruvate Sulfurtransferase) marks the path crossroad between low and high motile potential. Results from N-S dataset analysis. **c**. MPST overexpression in GB tissues compared to normal brain tissues. Expression in 163 glioblastoma (GB) versus 207 normal brain (Normal) tissues. Boxplot representation from GEPIA2 website. TCGA RNA-seq dataset. One-way ANOVA test, *: BH-adjusted p-value < 0.05. **d.** Schematic representation of the metabolic pathways over-mobilized in M^HIGH^ cells compared to M^LOW^ cells, as inferred from the KEGG pathway and trajectory modeling analyses.

Trajectory modeling was used next to delineate the acquisition of metabolic components potentially crucial for a motile cell state. Trajectory reconstruction based on expression of the motility signature genes was achieved with the STREAM python software using the N-S dataset which has the highest sequencing depth. The inferred trajectory corresponded to a branched path, highlighting an intermediary branch (S3-S1 branch) linking cells with low motile potential to cells with high motile potential (Fig.2b). Among the metabolic enzymes overexpressed in S3-S1 branch compared to each of the other branches, characterizing therefore the switch between low and high motile states (Supplementary table 5), two were overexpressed in M^HIGH^ cells of all datasets considered. *NIT2* encodes an omega-amidase involved in anaplerosis through shuttling of α-ketoglutarate and oxaloacetate from the glutamine and asparagine metabolisms into the TCA cycle (Cooper et al 2016). *MPST* (Mercaptopyruvate Sulfurtransferase) encodes an enzyme involved in cysteine metabolism and is part of anti-oxidative cell defenses. MPST converts 3-mercaptopyruvate, derived from cysteine transamination, to pyruvate while transferring a sulfur to a thiophilic acceptor, thereby forming a persulfide (Filipovic et al 2018). MPST was selected for further studies based on the following rationale. MPST is overexpressed in GB tissues compared to normal brain tissues (Fig.2c). Mitochondria-located GOT2, an aminotransferase which ensures the formation of the MPST substrate 3-mercaptopyruvate, is overexpressed in GB cells with high motile potential (Supplementary table 2). MPST participates to anti-oxidant defenses, notably through persulfidation (also known as S-sulfhydration) of proteins (P-SSH), a post-translational modification of reactive cysteine residues that protects proteins from irreversible cysteine hyperoxidation (Filipovic et al 2018) (Pedre & Dick 2021). Persulfidated MPST (MPST-SSH) has also been proposed as a key intermediate in production of GSH-persulfides (GSSH) (Kimura et al 2017). Although MPST participation in the control of GB cell properties is unknown, evidences have been provided for its stimulatory effect on the migration of endothelial cells (Tao et al 2017) (Abdollahi Govar et al 2020) and murine colon cancer cells (Augsburger et al 2020). Taken with the results from our in silico analyses, these data led us to postulate that enhanced oxidative stress coupled with MPST mobilization plays a key role in GB cell motility (Fig.2d).

### Enhanced ROS production in cells with high motile potential

The predictive values of the results from the computational analyses were experimentally evaluated using GB patient-derived cells (PDC). PDC relative migratory and invasive properties, as determined using spheroid-on-Matrigel migration assays, invasion-across-collagen assays and Matrigel-coated transwell assays showed 5706**-PDC to be the most motile, followed by R633- and P3-PDC (Supplementary Figure 4A-C).

We first determined whether Reactive Oxygen Species (ROS) affect GB cell motility. Intracellular ROS levels were evaluated with CellROX Green or Deep red fluorogenic probes. Cell exposure to the anti-oxidant N-Acetyl Cysteine (NAC) or the ROS generator menadione resulted in decreased or increased fluorescent signals detected by FACS, thereby confirming CellROX suitability for detecting ROS in GB PDC (Fig. 3a). Microscopic observation of PDC-spheroids first seeded on Matrigel and then labeled with CellROX Green probe showed a higher ROS signal in GB cells migrating out of the spheroids compared to their static counterparts (Fig.3b).

**Fig. 3.**
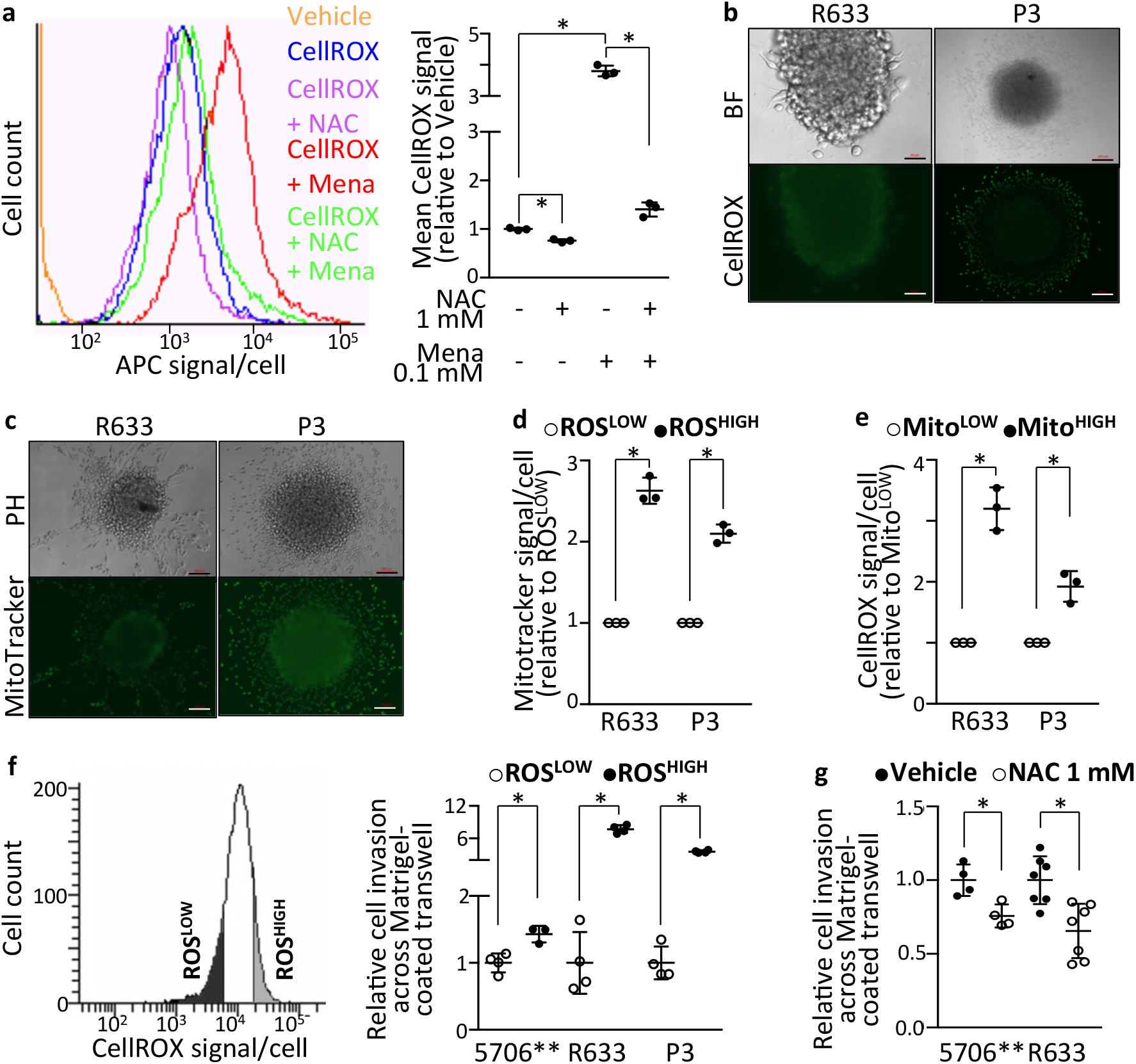
Motile glioblastoma cells exhibit enhanced ROS production and higher mitochondrial mass than their static counterparts. **a.** FACS analysis of CellROX signal in basal conditions, and following cell treatment with the anti-oxidant N-Acetyl Cysteine (NAC, 1mM, 1h) or the ROS generator menadione (Mena, 0.1mM, 30min). 5706** PDC. Note the decrease in fluorescent signal in NAC-treated PDC and the increased fluorescent signal in Menadione-treated PDC. Mean ± SD, n = 3 independent biological samples, *: p < 0.05, unpaired t-test with Welch’s correction. **b.** Higher ROS production detected in cells migrating out of spheroids. ROS levels assessed using CellROX Green reagent. R633 and P3 PDC. Brightfield (top panel) and CellROX fluorescence signal (488nm, bottom panel) imaging. Scale bars = 50μm (R633) and 200μm (P3). **c.** Microscopic visualization of mitochondrial mass using MitoTracker Green reagent. Scale bars = 200μm. **d.** Higher mitochondrial mass in cells with high ROS production compared to cells with low ROS production. R633 and P3 PDC. FACS analysis of ROS levels using CellROX Deep Red reagent and mitochondrial mass using MitoTracker Green reagent. Cells classified based on their basal ROS levels. Mean ± SD, n = 3 independent biological samples, *: p < 0.05, unpaired t-test with Welch’s correction. **e.** Cells with high mitochondrial mass have higher ROS production than cells with low mitochondrial mass. R633 and P3 PDC. FACS analysis of ROS levels using CellROX Deep Red reagent and mitochondrial mass using MitoTracker Green reagent. Cells classified based on their mitochondrial mass. Mean ± SD, n = 3 independent biological samples, *: p < 0.05, unpaired t-test with Welch’s correction. **f.** Cells with high ROS production have higher invasive properties than cells with low ROS production. 5706**, R633 and P3 PDC. Left panel: Example of FACS-sorting of PDC into ROS^LOW^ and ROS^HIGH^ fractions. Right panel: Cell invasion across Matrigel-coated transwells. Mean ± SD, n = 3-4 independent biological samples, *: p < 0.05, unpaired t-test with Welch’s correction. **g.** Decreasing ROS levels in GB-PDC using 1mM NAC decreases cell invasion. 5706** and R633 PDC. Cell invasion across Matrigel-coated transwells. Mean ± SD, n = 4-7 independent biological samples, *: p < 0.05, unpaired t-test with Welch’s correction.

Considering this result with our prior observation of enriched expression in genes involved in the TCA cycle and the ETC, we measured mitochondrial mass using the MitoTracker Green reagent. An increased mitochondrial load was detected in cells migrating out from cell spheroids (Fig. 3c). FACS was then used to isolate PDC according to their CellROX or Mitrotracker signals. Cells with high ROS production exhibited a 2.1-2.6 times higher mitochondrial mass than cells with low ROS production (Fig.3d). Conversely, cells with high mitochondrial mass exhibited higher ROS production than cells with low mitochondrial mass (1.9-3.2 times more) (Fig.3e). We next assessed the invasive potential of GB PDC with high ROS production. To do so, we FACS-sorted GB PDC according to their basal ROS production, as detected with CellROX probe (Fig.3f). Evaluation of the ability of each cell population to invade through Matrigel-coated transwells showed a 1.4-7.7-fold increase in invasiveness of GB cells with high ROS levels compared to cells with low ROS levels (Fig.3f). Finally, we examined the effects of decreasing ROS levels on cell motility by exposing the most motile PDC (5706** and R633) to NAC. Decreasing ROS levels with NAC treatment resulted in decreased GB cells’ ability to invade through Matrigel-coated transwells (Fig.3g). These results indicate that motile GB cells exhibit enhanced ROS production, as predicted by the results of the computational analyses, and that ROS participate to GB cell motility.

### MPST enzymatic activity is required for GB cell motility

To determine if MPST is necessary for GB cell motility, we first assessed MPST protein levels in motile GB cells by immunocytochemistry. We observed higher MPST protein levels in cells that migrated out of spheroids or GBO (Fig. 4a). We next knocked down *MPST* expression in the three GB PDC with high and intermediate motile potential, using lentiviral transduction of small hairpin (sh) RNA. Cell transduction with shMPST lentiviral constructs led to decreased MPST mRNA and protein levels with no major cytotoxic effect (Supplementary Figure 4D-F) Similar results were obtained with another shMPST construct (Supplementary Figure 4G-I). Knocking down *MPST* expression resulted in a sharp decrease in cell migration (51-86%) as determined with spheroid-on-Matrigel assays (Fig.4b). Monitoring cells’ ability to move across micro-channels with periodic 3μm mechanical constraints in a microfluidic chip further showed that *MPST* knockdown also impaired GB cells’ ability to change their shape so as to migrate through narrow and tortuous spaces (Fig.4c). Cell invasion was also decreased by 44-95% upon *MPST* knockdown as observed using spheroid-in-collagen (Fig. 4d) and Matrigel-coated transwells invasion assays (Fig.4e). The observed decrease in cell motility can be due to an inhibition of MPST catalytic activity or to defects in other MPST functions independent from its enzymatic activity. To distinguish between these two possibilities, we treated cells or GBOs with 200 μM I3-MT-3, a selective pharmacological inhibitor of MPST enzymatic activity (Hanaoka et al 2017). Spheroid-on-Matrigel assays showed that I3-MT-3 decreased PDC migration compared to vehicle-treated cells (Fig.4f). Alike PDC, cell migration out of GBOs was also inhibited by I3-MT-3 (Fig.4f). I3-MT-3 inhibited also PDC migration through constraining microfluidic chip channels (Fig. 4g). These results indicated that MPST catalytic activity was required for GB cell motility. Given that MPST participates to the cells’ anti-oxidative defenses through its role in protein persulfidation, we further determined protein persulfidation levels in GB PDC with decreased MPST expression or activity using the recently published Dimedone-Switch method (Zivanovic et al 2019). Decrease in the global protein persulfidation levels upon *MPST* knockdown (Fig.5a), and upon MPST enzymatic inhibition (Fig.5b), showed that MPST participates to protein persulfidation in GB cells.

**Fig. 4.**
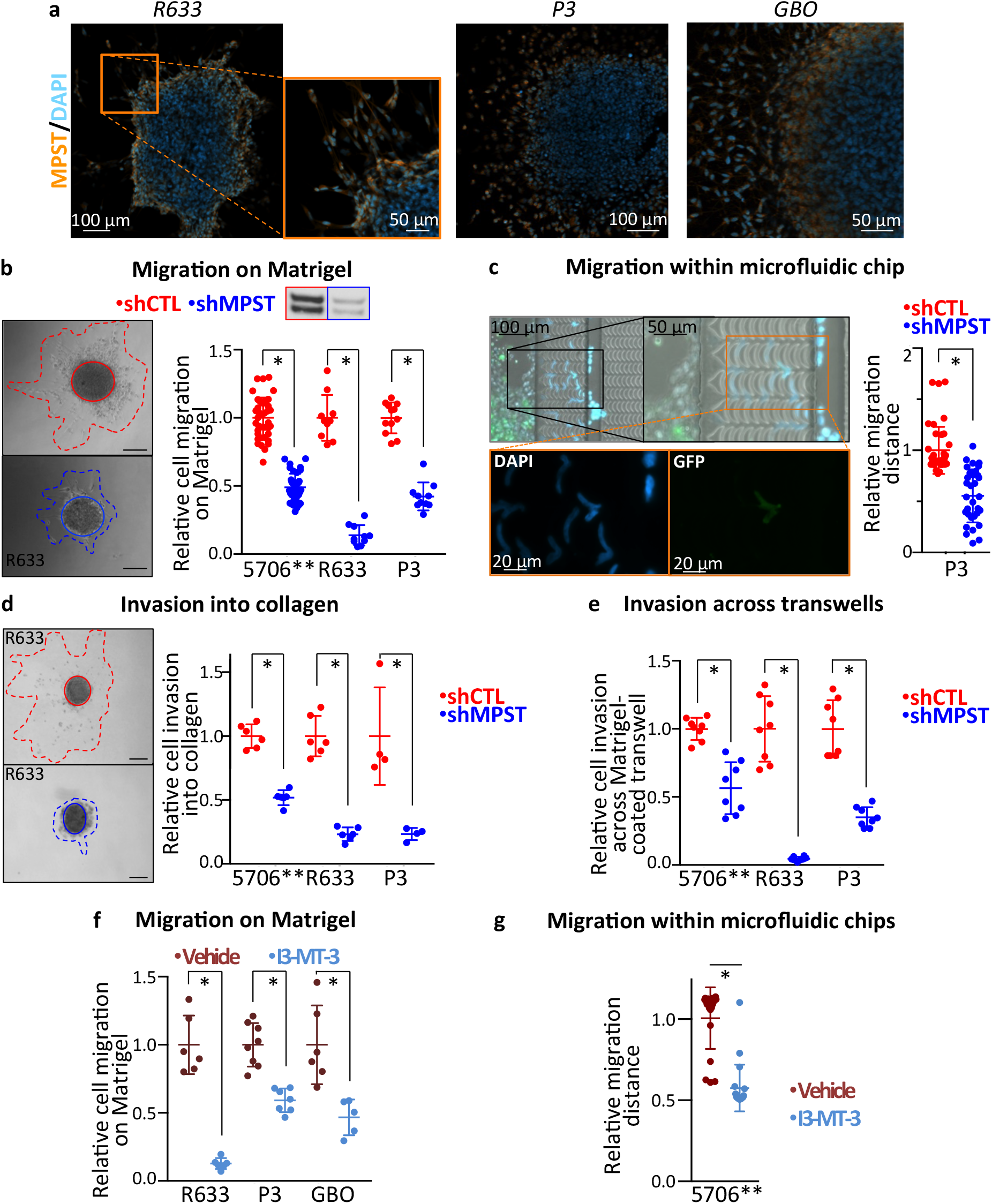
MPST is required for glioblastoma cell migration and invasion. **a.** MPST immunolabeling of GB cells migrating out of PDC spheroids or GBOs. MPST signal in red. Nuclei DAPI staining in blue. **b.** *MPST* knockdown decreases cell migration on Matrigel. 5706**, R633 and P3 PDC. Microphotographs: examples of cell migration assays, scale bars = 200μm, solid line = spheroid core, dotted line = migration area. Graph: quantification of the migration 7h (5706**) or 24h (R633 and P3) post-seeding. Mean ± SD, n = 10-41 independent biological samples, Mann-Whitney test, *: p < 0.05. Inset above graph illustrates decreased MPST protein levels (MW: 33/35 kDa) in PDC expressing *shMPST.* Western Blot analysis. **c.** *MPST* knockdown decreases cell migration into microfluidics chips. PDC expressing shMPST and GFP, were loaded together with equal numbers of shControl-PDC. Only rare shMPST-PDC (green cells) cross the microchip channels, contrary to shControl-PDC (white cells). Scale bar = 200μm. Graph: distances travelled by the cells over 24hrs. Mean ± SD, n = 32-40 from 4 independent biological samples, *: p < 0.05, Mann-Whitney test. P3 PDC. **d.** *MPST* knockdown decreases cell invasion into collagen. Microphotographs: examples of invasion assays, scale bars = 200μm. Graph depicts the quantification of the invasion after 24h (R633- and P3-PDC) or 40h (5706**-PDC). Mean ± SD, n = 4-8 independent biological samples, *: p < 0.05, Mann-Whitney test. **e.** *MPST* knockdown decreases cell invasion across Matrigel-coated transwells. 5706**-, R633- and P3-PDC invasion assessed 24h post-seeding. Mean ± SD, n = 4-8 independent biological samples, *: p < 0.05, Mann-Whitney test. **f.** Inhibiting MPST enzymatic activity using the pharmacological inhibitor I3-MT-3 decreases cell migration. Cells treated with 200μM I3-MT-3 or vehicle for 72h (R633, P3) or 45h (GBO), and migration assessed 48h (R633, P3) and 45h (GBO) post-seeding on Matrigel. Mean ± SD, n = 5-8 independent biological samples, *: p < 0.05, Mann-Whitney test. **g**. Inhibiting MPST enzymatic activity with I3-MT-3 decreases cell migration into microfluidic chips. Graph: greatest distances travelled by the cells 24hr post-seeding. 5706**-PDC,. Mean ± SD, n = 20 from 2 independent biological samples,* p < 0.05, Mann-Whitney test.

Finally, we evaluated the consequences of *MPST* knockdown on overall tumor burden using orthotopic xenografts of PDC stably expressing luciferase and either shControl or shMPST. Bioluminescence imaging showed cell engraftment and initial tumor development whether the cells expressed shControl or shMPST, with delayed tumor formation observed for one cell line (Fig.5c). In all cases, we observed a reduced tumor burden in mice grafted with sh*MPST*-PDC, compared to mice grafted with shControl-PDC up to the experiment end-points (Fig.5d, Supplementary Figure 4J). Mice survival monitoring showed that mice xenografted with shMPST-PDC survived on average twice longer than mice xenografted with shControl-PDC (Fig.5e).

**Fig. 5.**
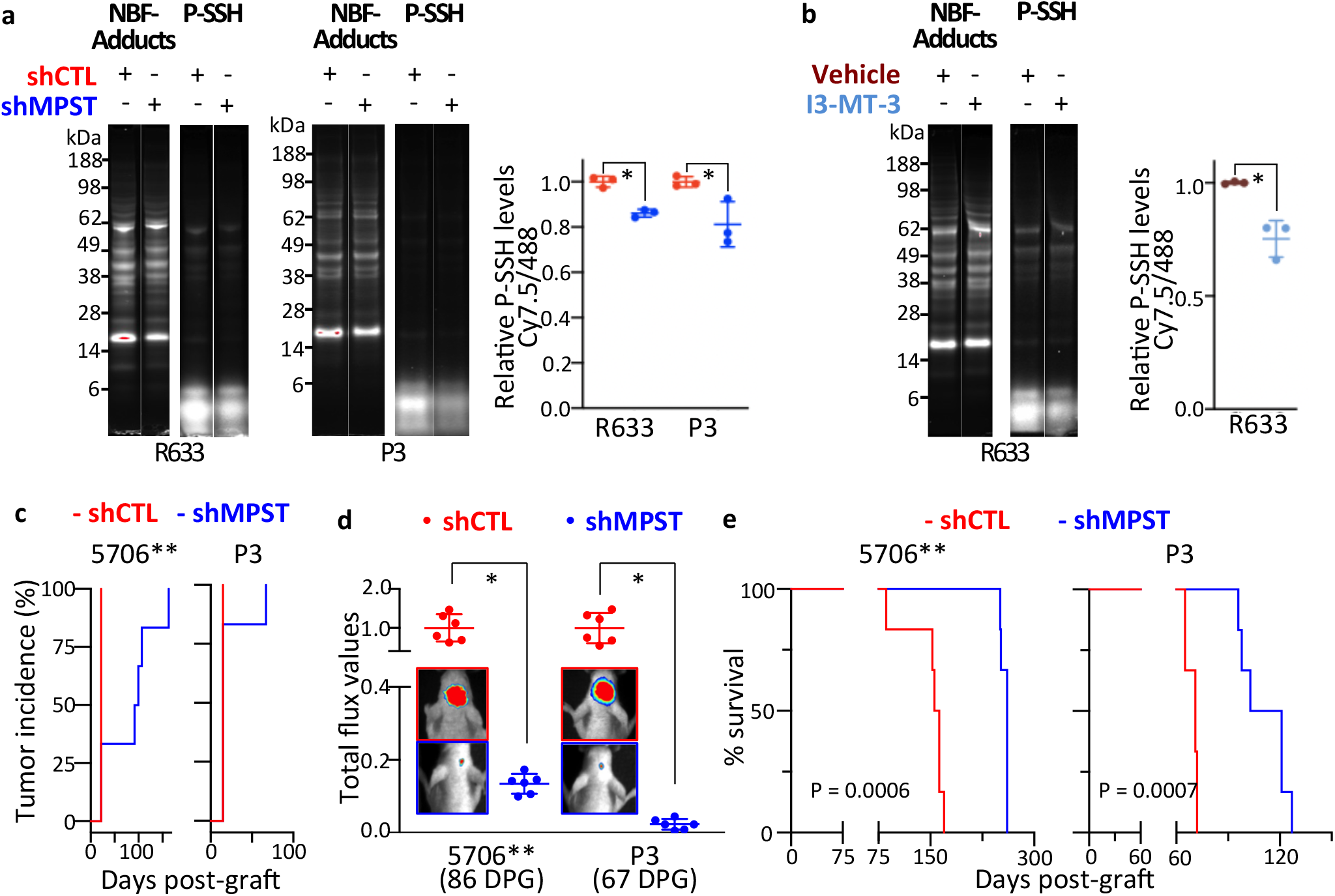
*MPST* knockdown decreases protein persulfidation, decreases tumor burden, and increases mice survival expectancy. **a.** Decreased protein persulfidation levels in sh*MPST*-PDC vs shControl-PDC. R633 and P3 PDC. In-gel detection of protein persulfidation (P-SSH) using the dimedone switch method with Cy7.5 as a reporting molecule and fluorescent signal of the NBF-adducts (488nm). P-SSH levels calculated as a ratio of Cy7.5/488 fluorescent signals. Mean ± SD, n= 3 independent biological samples, *: p < 0.05, unpaired t-test. **b.** Decreased protein persulfidation levels upon cell treatment with the MPST inhibitor I3-MT-3. R633 PDC. In-gel detection and calculation of P-SSH levels as in **i**. Mean ± SD, n= 3 independent biological samples, *: p < 0.05, unpaired t-test. **c.** *MPST* knockdown delays tumor development. Bioluminescent analyses of tumor growth initiated by grafting PDC transduced with a luciferase construct and either shControl (shCTL) or *shMPST* constructs. 5706** and P3 PDC. n = 6 mice per group. DPG: days post-graft. **d.** *MPST* knockdown decreases tumor burden as shown by quantification of the tumor bioluminescent signals. 5706** and P3 PDC. DPG: days post-graft. Mean ± SD, n = 6 mice per group, *: p < 0.01, Mann-Whitney test. **e.** Kaplan-Meier survival curves demonstrating a significant survival benefit of mice grafted with PDC expressing *shMPST* compared to mice grafted with PDC expressing shControl. 5706** and P3 PDC. n = 6 mice per group. Log-rank (Mantel-Cox) test.

## Discussion

The diversity of cancer cells populating GB is a major challenge in treating these brain cancers. Understanding the molecular basis of the varying functional cell states that co-exist within GB at the time of patients’ diagnosis and are commonly shared across tumors, opens a path to overcome this challenge. A way to access such information is coupling experimental validations with computational modeling of transcriptomes of single cells sorted from patient GB, which offer the closest possible map of the patients’ tumor cells. Following this blueprint led to circumscribing the overall metabolic characteristics of motile cells across thirty patients’ GB, and to identifying MPST as a metabolic enzyme crucial for GB cell motility and tumor development.

Our results suggest that GB cells with high motile potential are characterized by greater energetic production than cells with low motile potential, as shown by the enrichment in multiple energetic pathways. This is in agreement with motile cells needing more energy to colonize surrounding brain tissue and adapt to new microenvironments with unique nutrient and oxygen availability, thereby mobilizing different pathways to meet their high energetic needs (Garcia et al 2021). Of note, this overall energetic mobilization appears to converge towards the TCA and the ETC, a modeling result supported by the experimental demonstration of enhanced ROS production and higher mitochondrial mass load in motile GB cells. These results are in coherence with previous studies on epithelial cancer cells showing that an ETC overload with preserved mitochondrial functions and an increased mitochondrial superoxide production promote tumor cell migration and invasion (Porporato et al 2014). Results from previous studies, using GB cell lines cultured in serum, highlight also a role for mitochondria in cell motility. An example is the trafficking of energetically active mitochondria to membrane protrusions accompanying enhanced invasive properties of the LN229 GB cell line (Caino et al 2015). Another is the inhibitory effect of ROS scavenging on EGF-stimulation of T98G cell migration (Pudelek et al 2020). More broadly, ROS are known to promote activity of pro-migratory signaling pathways by direct redox modification of their protein components (Weng et al 2018) (Dustin et al 2020).

Intracellular ROS levels must however be tightly controlled since ROS overload can have severe deleterious consequences on cancer as well as on normal cell properties and viability. Coherently, our results indicate that GB cells with high motile potential are enriched in anti-oxidant pathways capable of counteracting oxidative stress. We observed an upregulation of the pentose phosphate pathway (PPP) that implicates enhanced NADPH production necessary for the function of several antioxidant proteins. We also observed enhanced glycine, glutamate and cysteine metabolism together with enhanced glutathione metabolism that implicates enhanced availability of the major anti-oxidant compound GSH. Cysteine metabolism involvement in the anti-oxidative process extends beyond supporting GSH synthesis. The CSE (Cystathionine Gamma-Lyase also known as CGL, encoded by *CTH),* CBS (Cystathionine Beta-Synthase) and MPST (also known as MST) enzymes of the cysteine metabolism participate to protein protection from detrimental hyperoxidation of their reactive cysteine residues (Zivanovic et al 2019).

MPST was the only metabolic enzyme participating in protection from oxidative stress predicted by trajectory modeling to be at the crossroad of the path leading GB cells from low to high motility. Our results demonstrate the biological validity of this prediction. MPST knockdown resulted in a robust inhibition of GB cell motility. The cells exhibit a decreased ability to move around in a freely available space, as shown with spheroid-on-Matrigel migration assays and microfluidics assays. The latter shows in addition that the sharp reduction in the cells’ ability to migrate through narrow spaces is associated with impairment in cell deformability. MPST facilitation of cell migration might extend to cell types other than GB cells, MPST inhibition having previously been involved in the migration of a mouse carcinoma cell line (Augsburger et al 2020), and endothelial cells (Tao et al 2017) (Abdollahi Govar et al 2020). We also observed a robust decrease in GB cell invasiveness upon MPST knockdown, GB cells exhibiting an impaired ability to remodel the ECM so that they can move into their surroundings as shown with spheroid-in-collagen and Matrigel-coated transwell assays. In coherence with these *in vitro* results, alteration of GB cell properties by MPST knockdown translates *in vivo* into reduced tumor burden, and a robust increase in mice survival, albeit the cells retain their tumor-initiating properties. MPST enzymatic activity is required for its pro-motile effects, the MPST pharmacological inhibitor I3-MT-3 (Hanaoka et al 2017) having the same consequences as MPST knockdown on GB cell migration. MPST catalyzes stepwise the desulfuration of 3-mercaptopyruvate that generates an enzyme-bound persulfide, and the transfer of the persulfide’s outer sulfur atom to proteins or small molecule acceptors (Pedre & Dick 2021). MPST knockdown was accompanied with reduced persulfidated protein levels in GB cells, indicating that protein persulfidation is part of the mechanisms by which MPST affects GB cell motility. Whether the advantage conferred by this post-translational modification stems from a mere overall protection of proteins from hyperoxidative damage or from more specific consequences remains to be determined. The few surveys of persulfidated proteins performed without exogenous dihydrogene sulfide (H_2_S) supply showed that persulfidation affected proteins involved notably in energetic pathways (Ida et al 2014) (Gao et al 2015) and migration and associated signaling pathways (Zivanovic et al 2019) (Murphy et al 2019) (Zuhra et al 2021). For instance, proteins of the glycolysis pathway were reported to be persulfidated in a human pancreatic beta cell line (Gao et al 2015). The resulting increase in glycolytic metabolic flux could stem from enhanced catalytic activity of glycolytic enzymes such as GAPDH or LDHA under their persulfidated form (Mustafa et al 2009) (Gao et al 2015) (Untereiner et al 2017). Persulfidation studied in HEK293 cells has also been reported to enhance actin polymerization and alter actin-dependent cytoskeletal rearrangements (Mustafa et al 2009). Interestingly, several proteins pertaining to migration, including the focal protein kinase encoded by PTKN2, have also been found to undergo persulfidation in HeLa cells treated with EGF, a known pro-migratory signaling factor (Zivanovic et al 2019). Most of these increases in protein persulfidation are presumably mediated by CBS and/or CSE activity, as supported by studies implementing inactivation of these enzymes and/or by their conspicuous expression in the biological systems under scrutiny (Mustafa et al 2009) (Gao et al 2015) (Untereiner et al 2017) (Zivanovic et al 2019) (Zuhra et al 2021). MPST-mediated protein persulfidation has so far been reported for only two proteins, the sulfurtransferase Mocs3 and thioredoxin (Trx). Through Mocs3, it is involved in a cascade leading to protein urmylation and tRNA thiolation, and through Trx in H_2_S generation (Pedre & Dick 2021). The overall decrease in protein persulfidation levels we observed in GB cells upon MPST knockdown indicates that MPST affects a wider range of proteins.

Sulfur metabolism, considered from the perspective of H_2_S levels detected in tumor tissue homogenates, has been reported to be lower in GB than in normal brain tissues, and controlled H_2_S production appears to be necessary for tumor fitness as increasing H_2_S levels with H_2_S donors decreases GB cell viability, and development of subcutaneous mouse GB cell grafts (Silver et al 2021). Finding decreased H_2_S production upon inhibition of CBS expression or activity in mouse GB cells (Silver et al 2021), and in a serum-cultured cell line (Takano et al 2014), point to CBS as the mediator of H_2_S production in GB as in other cancer cells studied so far. In this context, our results unravel a novel role for sulfur metabolism in the control of GB development through MPST. Demonstration of the participation of MPST in the control of GB cell motility was made possible by taking into account GB heterogeneous populations through the consideration of the cell functional state in link with its metabolic underliers. Only single cell data allow analysis at this level of granularity, avoiding confusion brought up by averaging different cell types and functional states in a whole tissue or cell culture. However, it remains unknown whether our modeling results pertain to a specific mode of cell motility and/or path of migration. Both individual and collective migrations have been identified in GB (Vollmann-Zwerenz et al 2020) (Volovetz et al 2020). Enrichment of oRG-like gene modules in M^HIGH^ cells suggests that the motile cells we captured move in a solitary fashion like oRGs in the developing cortex. However, it is yet to be demonstrated that gene markers for oRG-like GB cells are indeed supporting individual migration rather than collective migration. With respect to the path of migration, we found that MPST knockdown impaired cell invasiveness not only through collagen, a major component of the tumor ECM, but also through Matrigel, whose composition mixes its major laminin component to collagen IV and heparan sulfate proteoglycans, similar to the blood vessels’ basement membrane found in GB (Ljubimova et al 2006). These results suggest that MPST could be necessary for GB cells moving along blood vessels as well as through the brain parenchyma. In the different datasets considered in this study, the edge or core origin of the cells is most often unknown, though they are most likely coming from dense tumor cell regions. Removal of the surrounding parenchyma is usually undertaken with great care by neurosurgeons in order to spare at the best motor and cognitive functions. Although Darmanis and colleagues distinguished malignant cells isolated from the tumor core and the periphery, the cancer cells isolated from the latter are too few to authorize robust computational analyses. It is therefore very likely that our findings relate to dense tumor areas’ cells, moving either along the blood vessel network that characterize these highly angiogenic tumors, or across the tumor parenchyma enriched in components favoring cell displacement.

## Methods

Schemes were created with BioRender.com. All bioinformatics analyses were performed using the R software version ≥ 3.6.1 (https://cran.r-project.org/) or Python 3.7.9 (https://www.python.org/). All resources and materials, R packages, corresponding websites and references are listed in Supplementary Table 6.

### scRNA-seq data pre-processing

Four publicly available single-cell transcriptomic datasets from GB patient tumors were downloaded, including the only two publicly-available datasets obtained with the SMART-seq2 technique (Neftel et al 2019) (Darmanis et al 2017), and two datasets obtained with the 10X Genomics technique (Neftel et al 2019) (Pombo Antunes et al 2021) (Supplementary Table 6). The last two datasets were selected among possible others because of the availability of the raw UMI counts. We designated them as NS, D-S, N-10X and PA-10X, respectively. Outlier cells with low-complexity transcriptomes, revealed by plotting the graphical distribution of the number of detected transcripts and genes per cell, were excluded from the analyses as described (Saurty-Seerunghen, 2019). For 10X datasets, detection of over 20% mitochondrial genes per cell was considered as indicative of dying cells leading to their exclusion from the analyses. Therefore, we retained for further analyses 4916 malignant cells from twenty adult GB patients in the N-S dataset, 1033 malignant cells from four adult GB patients in the D-S dataset, 5797 malignant cells from six adult GB patients in the N-10X dataset and 8666 malignant cells from six adult GB patients in the PA-10X dataset. Log_2_-transformed Counts Per Million (log_2_(CPM+1)) were used for D-S and the 10X datasets, unless otherwise specified. CPM corresponds to the counts of gene-mapped reads normalized by the total number of mapped reads per cell divided by one million, thus allowing comparison of read abundance across libraries of different sizes. For the N-S dataset, gene expression data currently available are in log_2_((TPM/10) + 1). TPM (Transcripts Per Million) normalization takes into account an eventual biased estimation of long transcript numbers, by dividing the number of mapped reads by the transcript’s length. Since expression values in CPM could not be calculated from these data, log_2_((TPM/10) + 1) values were just transformed into log_2_(TPM + 1) to optimize the comparison between the different datasets. To avoid potential analytical bias due to scarcely detected genes, genes detected in less than 1% cells were filtered out.

Malignant and normal cells were distinguished either according to the cell annotations provided for both SMART-seq2 datasets, or when absent as for 10X Genomics datasets, based on inference of copy-number variations (CNVs), a hallmark of malignant cells. Expression data in log_2_(CPM/100 + 1) values were used, which shrinks expression ranges, allowing to keep only the major gene expression variations, based on which malignant and normal cells can be distinguished. Data were processed using a three-step approach: CNV inference, marker gene expression and unsupervised cell clustering. CONICSmat R package used to infer CNVs has the advantage of not requiring an a priori reference cell dataset (Muller et al 2018). The default filtering and normalization procedures were followed, as outlined in https://github.com/diazlab/CONICS/wiki/Tutorial--CONICSmat;--Dataset:-SmartSeq2-scRNA-seq-of-Oligodendroglioma. CONICSmat fits a two-component Gaussian Mixture Model (GMM) to the average gene expression across all cells within each chromosomal region. As a result, genes of a given region present a lower expression in cells with a deletion of this region than cells without the deletion. The opposite is observed for genes of amplified regions. The posterior probabilities of belonging to one of the two components of the model are then calculated for each cell. The copy number status across cells is predicted by the posterior probabilities for each cell belonging to the component with the higher mean. For each region, a CONICSmat likelihood ratio test adjusted p-value <0.001 and a difference in Bayesian Criterion >300 were retained as CNVs. The GMM-based CNV predictions were then used to group cells into potential malignant and non-malignant groups and were visualized using heatmaps (ComplexHeatmap R package (Gu et al 2016)) and UMAP (umap R package) plots based on 500 most variable genes (Supplementary Figure 5A-B, Supplementary Figure 6A-B). Second, expression of marker genes for pan-immune cells (PTPRC/CD45), macrophages (ITGAM, FCGR3A/CD16A, CD14), microglia (CSF1R, TMEM119), T-cells (CD2, CD3D) and oligodendrocytes (MOG, MAG) was highlighted on UMAP plots (Supplementary Figure 5C, Supplementary Figure 6C). Finally, a hierarchical clustering followed by a K-means clustering on the UMAP components (FactoMineR R package (Husson et al 2010)) was applied to identify cell groups that were most similar or different to one another (Supplementary Figure 5D, Supplementary Figure 6D). Using CNV status predictions, marker gene expressions and the clustering result, the cells were marked as malignant when they harbored CNVs, clustered together on UMAP and were devoid of normal cell markers (Supplementary Figure 5E, Supplementary Figure 6E).

R scripts used are provided in Supplementary Datafile 1.

### Cell grouping analyses

Clustering analysis was based on a molecular signature of ten elements (see results) and achieved using the Hierarchical Clustering on Principal Components (HCPC) approach (FactoMineR package (Husson et al 2010) modified to implement in a stepwise manner four methods for multivariate data analyses: Principal Component Analysis (PCA), Uniform Manifold Approximation and Projection (UMAP), hierarchical clustering, and partitioning clustering by the k-means (Supplementary Figure 1A). The ten principal components (PCs) identified by PCA were reduced non-linearly to two dimensions using UMAP. Euclidean distance was used to construct a cell-to-cell distance matrix based on the two UMAP components. Hierarchical clustering was then performed on this distance matrix using the Ward’s criterion (ward.D2 algorithm) in order to determine the number of clusters. The resulting partitioning of the cells was improved by a K-means clustering with ten iterations. The cell grouping was visualized using UMAP (umap R package) or chord plots (circlize R package (Gu et al 2014)). A Normalized Mutual Information (NMI) score (ClusterR package (Manning et al 2008)) was calculated to determine the contribution of cells issued from distinct tumors to each cluster, as described (Saurty-Seerunghen et al 2019). Of note, THBS1 was excluded from the motility signature when applied to PA-10X dataset due to its detection in <1% GB cells in that dataset.

Motility and other scores were obtained by computing the geometric mean of the expression values per cell of each element of the molecular signature corresponding to a given score. For each signature, only genes detected in ≥ 1% cells in ?3 datasets were considered. When null, expression values were imputed with a value of 1.

R scripts used are provided in Datafile S1.

### Analyses of genes differentially expressed between cell groups

Genes differentially expressed between cell groups with differing scores were identified using a Mann-Whitney (Wilcoxon Rank Sum) test with p-values adjusted for multiple testing (Benjamini-Hochberg (BH), adjusted p-value <0.01) (Soneson & Robinson 2018). Fold change (FC) was calculated as follows: FCi=xi-yi, where xi and yi are the log_2_ expression levels of gene i in conditions x and y, respectively. Only genes detected in at least 3% of GB cells were considered for this analysis. Genes coding for metabolism enzymes were identified using the list from KEGG (Kanehisa et al 2017).

Comparing list of differentially expressed genes between distinct datasets was performed with the venn R package on genes detected in ?3% malignant cells in all datasets, and after updating gene symbols based on gene metadata files downloaded from HGNC and NCBI websites (Supplementary Table 6). Genes absent from these metadata files, or with ambiguous symbols (e.g. genes whose current approved symbol is the previous symbol of another gene), or whose symbol was previously associated with more than one gene were excluded from the lists.

Gene ontology analyses were carried out on the enrichR website (Chen et al 2013) (Kuleshov et al 2016) (Xie et al 2021). We considered BH-adjusted p-value < 0.05 (Fisher’s Exact test) as the cut-off criterion for significance. Redundant terms were excluded. Graphs were generated using ggplot2 R package.

R scripts used are provided in Datafile S1.

### Trajectory inference analysis

To model the path taken by cells with low motile potential to reach a high motile potential, STREAM python package was used (Chen et al 2019). Briefly, the expression values of the ten elements of the motility signature were first extracted. Next, dimensions were reduced to four components using the spectral embedding algorithm (dimension_reduction STREAM function, other parameters are set to default). The components were then used for simultaneous tree structure learning and fitting using ElPiGraph (seed_elastic_principal_graph STREAM function, default parameters). The resulting trajectory structure was represented in a 2D subway map plot, where straight lines represent branches and each dot represents a single cell (plot_stream_sc STREAM function, dist_scale = 0.5 and root = ‘S5’, other parameters to default). To identify genes differentially expressed between cell populations from two adjacent branches, the detect_de_markers STREAM function was used. The mean of scaled gene expressions in each branch was calculated. Then, the fold change (FC) of mean expression between pairs of branches was computed, and Mann-Whitney U test performed. The U statistic was then standardized to Z-score to assess in which branch the gene is overexpressed. Genes with Z-score greater than 1 and log_2_FC greater than 0.15 are considered as overexpressed between two adjacent branches. Multiple testing correction was performed using the Benjamini-Hochberg method. Significance level was set at q value < 0.05.

### Biological material, lentiviral transduction and pharmacological treatment

Patient-derived cells (PDC) R633, 5706** and P3 obtained in our and other laboratories from neurosurgical biopsy samples of distinct primary GB were cultured in defined medium containing bFGF and EGF or bFGF only (P3), as described (El-Habr et al 2017) (Rosenberg et al 2017) (Eskilsson et al 2016). The PDC stably express luciferase, and 5706**-PDC express GFP as well. Cells were transduced with lentiviral vectors encoding a shRNA construct either neutral (shControl) or targeting *MPST* transcripts (shMPST) (Supplementary Table 6). Non-transduced cells were eliminated following a 1-2 week treatment with puromycin (1-2 μg/mL). The lentiviral particles were produced by the Plateforme vecteurs viraux et transfert de gènes (Necker Federative structure of research, University Paris Descartes, France). GBOs were generated and cultured from a primary GB patient tumor (Supplementary Table 6) following the method developed by Jacob and collaborators (Jacob et al 2020). In relevant experiments, cells were treated with 200μM I3-MT-3 or 1mM NAC (MedChemExpress and Sigma respectively) or their vehicles (DMSO and culture media respectively).

### Cell migration and invasion assays

Cell migration was assessed using spheroid-on-Matrigel assay, as described (Guyon et al 2020). Briefly, cells were counted as described (El-Habr et al 2017), and cell spheroids generated by seeding 2500 cells/100μL 0.4% methylcellulose (Sigma) in a U-bottom well (96 wells plate, Falcon). After a 24-48 hours culture, the spheres were seeded on Matrigel (0.2mg/mL culture media, Corning), and cell migration assessed after 7 hours (5706**) or 24 hours (R633, P3), unless specified otherwise. Images were acquired with a fluorescent microscope (Zeiss). The total area covered by GB cells migrating from a spheroid or GBO, as well as the central spheroid core area, were delineated and measured with FIJI software. The migratory index calculated corresponds to the ratio between the total area and the spheroid core area. Values were normalized to shControl or Vehicle. Microfluidic chips were designed and fabricated, as described (Courte et al 2018). Briefly, PDMS (Polydimethylsiloxane) blocks were modeled from a SU8 wafer encoding two 3mm-long 1mm wide and 50 μm-high cell culture chambers separated by 4.5μm-high arrays of archlike microstructures that encode successive mechanical constraints. The resulting SU8 wafers were used to cast polydimethylsiloxane (PDMS Sylgard, Ellsworth Adhesives) mixed at a 10:1 w:w ratio of base to curing agent. PDMS was cured 3 h minimum at 70 °C. The resulting chips were unmolded, and inlets were punched with a surgical biopsy punch (4 mm diameter) at both extremities of each compartment. The PDMS blocks were bonded to 130-160 μm thick glass coverslips (Fisher Scientific 11767065) after plasma surface treatment with an Atto plasma cleaner (Diener). Culture compartments were then immediately filled with deionized water. Devices were placed in individual Petri dishes for easier handling, and each of the resulting culture system was sterilized by exposure to a UV lamp for 30 min before use. 2*10^5^ cells PDC expressing shControl or shMPST were seeded at a 1:1 ratio into the cell reservoirs. The shMPST construct encoding GFP in addition to the shRNA allowed distinction between shControl and shMPST cells. Seven days post-seeding, cells were fixed with 4% paraformaldehyde, and nuclei stained using DAPI (Sigma). Cells were visualized with a fluorescent microscope (Zeiss). The distance travelled by the cells that moved away from the channel entrance was quantified using FIJI software. For the shMPST condition, only rare cells moved away from the channel entrance. We therefore considered all of them. For the shControl condition we considered an equivalent cell number among those having migrated the furthest away. For comparing I3-MT-3 effect on cell migration, 2*10^5^ vehicle- or I3-MT-3-treated cells were seeded in microfluidic chips. Cell migration was quantified by measuring the distance traveled by the ten cells having migrated the furthest away from the channel entrance.

Cell invasion was assessed using spheroid-in-collagen assay (Guyon et al 2020) and Matrigel-coated transwells (Renault-Mihara et al 2006). Spheroids generated as described above, were seeded in collagen (1mg/mL culture media, Corning), and cell invasion assessed 24 hours later. Images were acquired with a Zeiss microscope. The total area covered by GB cells leaving the spheroid and invading the collagen, as well as the central spheroid core area, were delineated and measured with FIJI software. The invasive index calculated corresponds to the ratio of the total area and the spheroid core area. Values were normalized to shControl or Vehicle. Transwells (8μm pores, Corning) were coated for 30min with Matrigel (0.2mg/mL culture media, Corning). They were then placed in wells containing culture media with growth factors. 40000 (R633 and P3) and 20000 (5706**) cells were seeded in 100μl media without growth factors into the transwells, and further cultured for 24 hours. Cells having invaded the bottom part of the membrane were counted after paraformaldehyde fixation and nuclei staining with DAPI (Sigma), as described (Renault-Mihara et al 2006).

### Detection of intracellular ROS levels

Cells or spheroids were incubated at 37°C for 30min with CellROX Green (5μM/R633, 10μM/P3) or Deep Red reagent (1.25μM/P3, 2.5μM/R633 and 5706**) (ThermoFisher, France). Fluorescence signal was then assessed by microscopy (CellROX Green, Zeiss microscope) or by FACS (CellROX Deep Red, ARIA II, BD Biosciences). When indicated, cells were treated with a ROS scavenger, N-Acetyl Cysteine (NAC, 1h, 37°C) and/or a ROS generator, Menadione (Mena, 30min, 37°C), and ROS levels were measured by FACS analysis. The latter was performed on 10000-gated events using Vdye-labeled cells for detecting any non-viable cell (1μL/4-5*10^5^ cells/mL, 30min, 4°C, eBiosciences). Violet laser (405nm) and Pacific Blue filter were used for Vdye detection, Blue laser (488nm) and FITC filter for CellROX Green reagent detection, and Red laser (640nm) and APC filter for CellROX Deep Red reagent detection. Data analysis and figure generation were performed using the FACS Diva software (BD Biosciences).

### Measure of mitochondrial mass

Cells, spheroids or GBO were incubated with 1μM MitoTracker Green reagent (Invitrogen) at 37°C for 30min and the fluorescence signal was assessed by microscopy or FACS. For FACS analysis, it was performed on 10000-gated events. Violet laser (405nm) and Pacific Blue filter were used for Vdye detection, and Blue laser (488nm) and FITC filter for MitoTracker Green reagent detection. Data analysis and figure generation were performed using the FACS Diva software (BD Biosciences).

### RT-QPCR assay of gene expression

Gene expression knockdown in response to shRNA expression was verified by RT-QPCR as previously described (Saurty-Seerunghen et al 2019) using the LightCycler480 (Roche, France) and the SYBR Green PCR Core Reagents kit (Bimake.com). The thermal cycling conditions comprised an initial denaturation step at 94°C for 5 min, and 40 cycles at 94°C for 30sec, 60°C for 30sec and 72°C for 30sec. Transcripts of the TBP gene encoding the TATA-box binding protein (a component of the DNA-binding protein complex TFIID) were quantified as an endogenous RNA control. Quantitative values were obtained from the cycle number (Cq value), according to the manufacturer’s manuals. Sequences of primers used for Q-PCR are given in Supplementary Table 6.

### Protein analyses

Cells were lysed using HEN Buffer (0.5M EDTA, 20% SDS, 1% NP-40, 10mM Neocuproine and 0.1M HEPES, pH 7.4) supplemented with 5mM 4-chloro-7-nitrobenzofurazan (NBF-Cl, Sigma-Aldrich), as described (Zivanovic et al 2019). Proteins were then precipitated twice in the presence of methanol and chloroform at 14000rpm for 15min at 4°C, the pellets being resuspended in 50mM HEPES (pH 7.4) supplemented with 2% SDS. Once completely dissolved, protein concentration was determined by BCA assay and adjusted to 3mg/mL. For detection of MPST protein levels, 30μg protein extracts were separated by SDS-PAGE using 4-12% (w/v) NuPAGE gels (Invitrogen, France) and transferred to Transblot nitrocellulose membranes (Biorad, USA). MPST detection was achieved using immunoblotting with anti-MPST (Sigma, 1:4000). The secondary antibody was anti-rabbit IgG (GE Healthcare, 1:30000). Signal detection was performed with SuperSignal™ West Femto Maximum Sensitivity Substrate chemiluminescence detection system (ThermoFisher, France). Densitometric analysis was achieved using ImageJ software. MPST levels were assessed by normalizing the chemiluminescent signal by the amount of proteins loaded assessed using Red Ponceau. Persulfidated protein levels were assessed following the Dimedone-Switch protocol, as described by Zivanovic and colleagues (Zivanovic et al 2019). Briefly, this method relies on the indiscriminate labeling of persulfides, thiols, sulfenic acids, and amino groups with 4-chloro-7-nitrobenzofurazan (NBF-Cl, the NBF-adducts yielding a fluorescent signal detected at 488nm). Upon incubation with dimedone-based probes, only the persulfides exchange their labeling, thus enabling their identification (red signal detected at 800nm). The protein extracts previously prepared in presence of NBF-Cl were incubated for 30min at 37°C with 25μM click mix provided by Dr Milos Filipovic (1mM Daz-2, 1mM Cy7.5 alkyne, 2mM TBTA Cu Complex, 4mM L-ascorbic acid, 30% acetonitrile and 20mM EDTA in PBS) and precipitated as described above. Proteins were subsequently resuspended in 50mM HEPES supplemented with 2% SDS and separated using SDS-PAGE on 4-12% (w/v) polyacrylamide gels. Gels were fixed in 12.5% methanol and 4% acetic acid for 30min. The Cy7.5 signal corresponding to persulfidated proteins was recorded at 800nm and the NBF-adducts signal was recorded at 488nm on a Chemidoc Imager (BioRad, France). Persulfidation levels were assessed by normalizing the Cy7.5 intensity to Alexa488 (NBF-adducts) intensity.

### Immunocytochemistry

Cells were harvested, PBS washed, smeared on SuperFrost slides (ThermoScientific, France), and fixed with 4% paraformaldehyde at room temperature. Following fixation, cells were washed with PBS, and incubated for 60 min at room temperature in Antibody blocker/diluent (Enzo). MPST primary antibody (Sigma, 1:50) was incubated overnight at 4°C. Secondary antibody anti-rabbit A555 was incubated at room temperature for 1 h (1:500, Life Technologies). Nuclei were stained using DAPI (Sigma). Immunostaining was analyzed with a fluorescent microscope equipped with an ApoTome module (Zeiss). Immunofluorescent signals were analyzed with Zen software (Zeiss) using 2.5 μm optical sections.

### Intracerebral xenografts

The animal maintenance, handling, surveillance and experimentation were performed in accordance with and approval from the Comité d’éthique en expérimentation animale Charles Darwin N°5 (Protocol #5379). 5706** and P3 PDC transduced with lentiviruses encoding either a shControl or a shMPST, were used. 1.5*10^5^ cells were injected stereotaxically into the striatum of anesthetized 8-10 weeks-old Nude female mice (Janvier Laboratories, France), using the following coordinates: 0mm posterior and 2.5mm lateral to the bregma, and 3mm deep with respect to the surface of the skull. Tumor formation and development were monitored by bioluminescence imaging performed on a Photon Imager Biospace (Biospace Lab, France), after intraperitoneal injection of 150μL luciferin (20mM, ThermoFisher, France). Bioluminescent signals were visualized with M3 Vision software (Biospace Lab, France).

### Statistical analyses

R version 3.6.1 or Prism 7.0 software (GraphPad) were used to generate plots and for statistical analyses. Significance level was set at p < 0.05, unless otherwise indicated. The type of statistical test used is provided in the figure legends. All experiments were performed using independent biological samples. Mean ± SD are shown.

## Acknowledgements

We are deeply grateful to Dr Milos Filipovic and Thibaut Vignane (SULFAGING group, ISAS, Dortmund, Germany) for their gift of reagents and for sharing protocols for assessing persulfidated protein levels. All our renewed thanks to Dr François-Xavier Lejeune (ICM, Salpétrière Hospital, Paris) for his advice on statistical analyses. We acknowledge the contribution of AniRA lentivectors production facility from the CELPHEDIA Infrastructure and SFR Biosciences (UAR 3444/CNRS, US8/Inserm, ENS de Lyon, UCBL), especially Gisèle Froment, Didier Nègre and Caroline Costa. This work was supported by grants from Région Ile-de-France (MSS fellowship), INCADGOS-Inserm_12560: SiRIC CURAMUS (financially supported by the French National Cancer Institute, the French Ministry of Solidarity and Health and Inserm), and La Fondation pour la Recherche Médicale – Equipes FRM 2020.

## Author Contributions

Conceptualization, MPJ, EEH, TD, HC, MSS; Methodology, MSS, LB, CA, TD, AI, JMP, TV, EEH and MPJ; Experimental investigations, MSS, VD, GC, JG, AR, SF, FA, FBV, LT, MPJ; Computational analyses, MSS and LB; Writing – Original Draft, MSS and MPJ; Writing – Review & Editing, all; Supervision, MPJ and EEH; Project Administration, HC; Funding Acquisition, HC, MPJ.

## Supplementary Figure legends

**Supplementary Figure 1, related to Fig.1.**
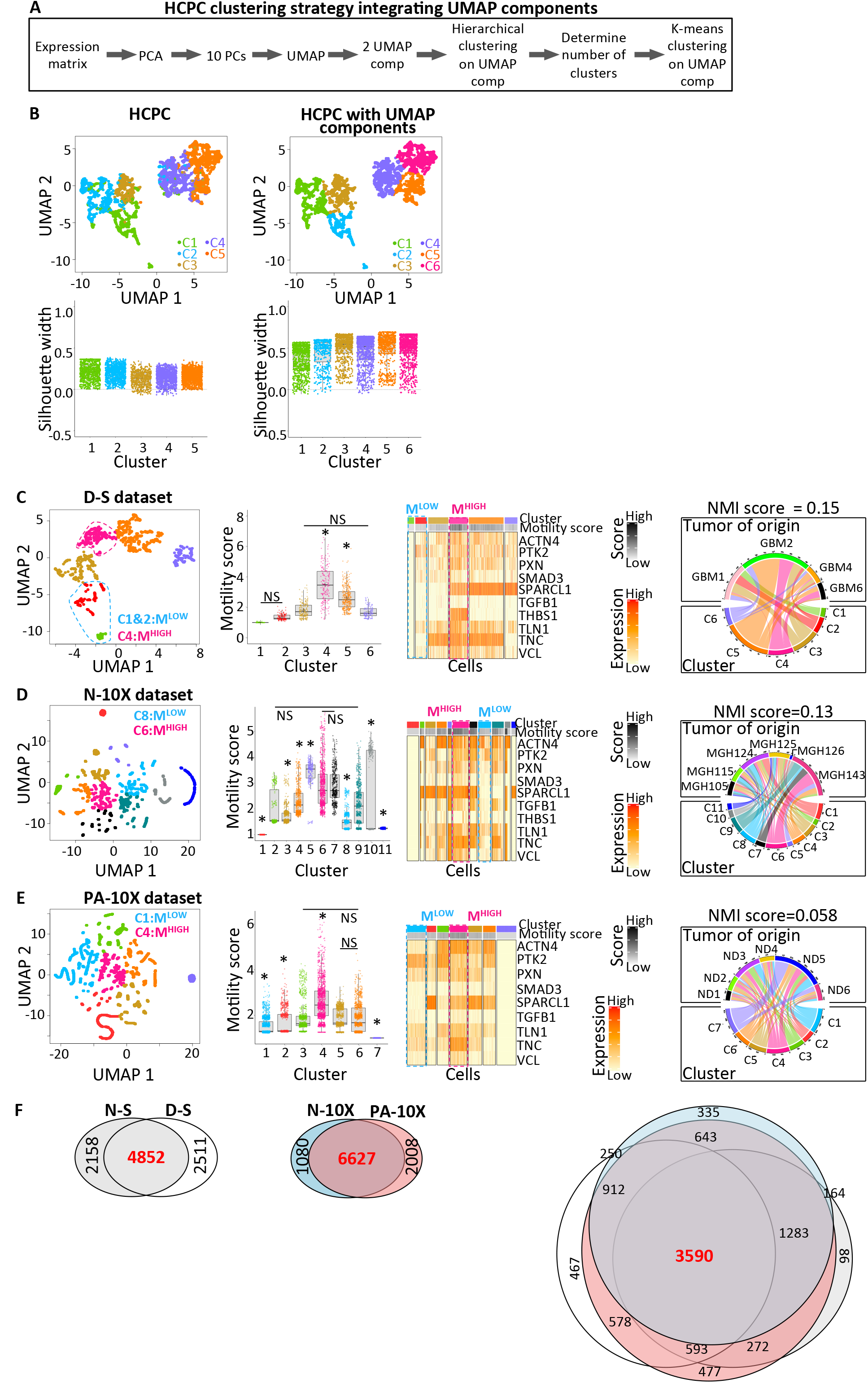
**A-B. Grouping strategy A**. Schematic representation of HCPC clustering strategy integrating UMAP components. comp: components, HCPC: Hierarchical clustering on principal components, PCA: Principal component analysis, PC: Principal components, UMAP: Uniform Manifold Approximation and Projection. **B**. Integrating UMAP components results in more homogeneous clusters, as shown by elimination of cell outliers on the UMAP representations (compare upper left and right panels) and the increase in the Silhouette width index (compare lower left and right panels). Clustering based on expression of the ten elements of the motility signature, and performed using 4916 glioblastoma cells from N-S dataset. **C-F**. **Motility signature captures cells with similar profiles in independent datasets.** Clustering malignant cells from D-S (**C**), N-10X (**D**) and PA-10X (**E**) datasets based on the motility signature genes. From left to right subpanels: UMAP representation; identification of cell groups with the highest and lowest mean motility scores (M^HIGH^, M^LOW^); Heatmap representation of the relative expression of each signature element per cluster, Contribution of each tumor to identified clusters. *: Clusters with mean motility score statistically different from each of the other clusters, p < 0.01, one-way ANOVA, Tukey’s multiple comparisons test, NS: non-significant. **F**. Overlaps between lists of genes overexpressed (OEG) in M^HIGH^ compared to M^LOW^ cell groups from distinct datasets. High overlap between lists from datasets obtained with the same sequencing techniques, with 69.2% of OEG in M^HIGH^ cells from N-S dataset also overexpressed in M^HIGH^ cells from D-S dataset (4852/7010), and 86% of OEG in M^HIGH^ cells from N-10X dataset also overexpressed in M^HIGH^ cells from PA-10X dataset (6627/7707). When comparing N-S, D-S, N-10X and PA-10X to the 3 other datasets, lists of OEG in M^HIGH^ cells overlap by 56.6% (3590/6348), 53.2% (3590/6750), 48.4% (3590/7418) and 43.0% (3590/8348), respectively. Due to graphical constraints, two overlaps are not shown on the third venn diagram: 119 genes were identified as overexpressed in M^HIGH^ cells in the two SMART-seq2 datasets only, and 241 overexpressed in M^HIGH^ cells from N-S, D-S and N-10X datasets (not from PA-10X).

**Supplementary Figure 2, related to Fig.1.**
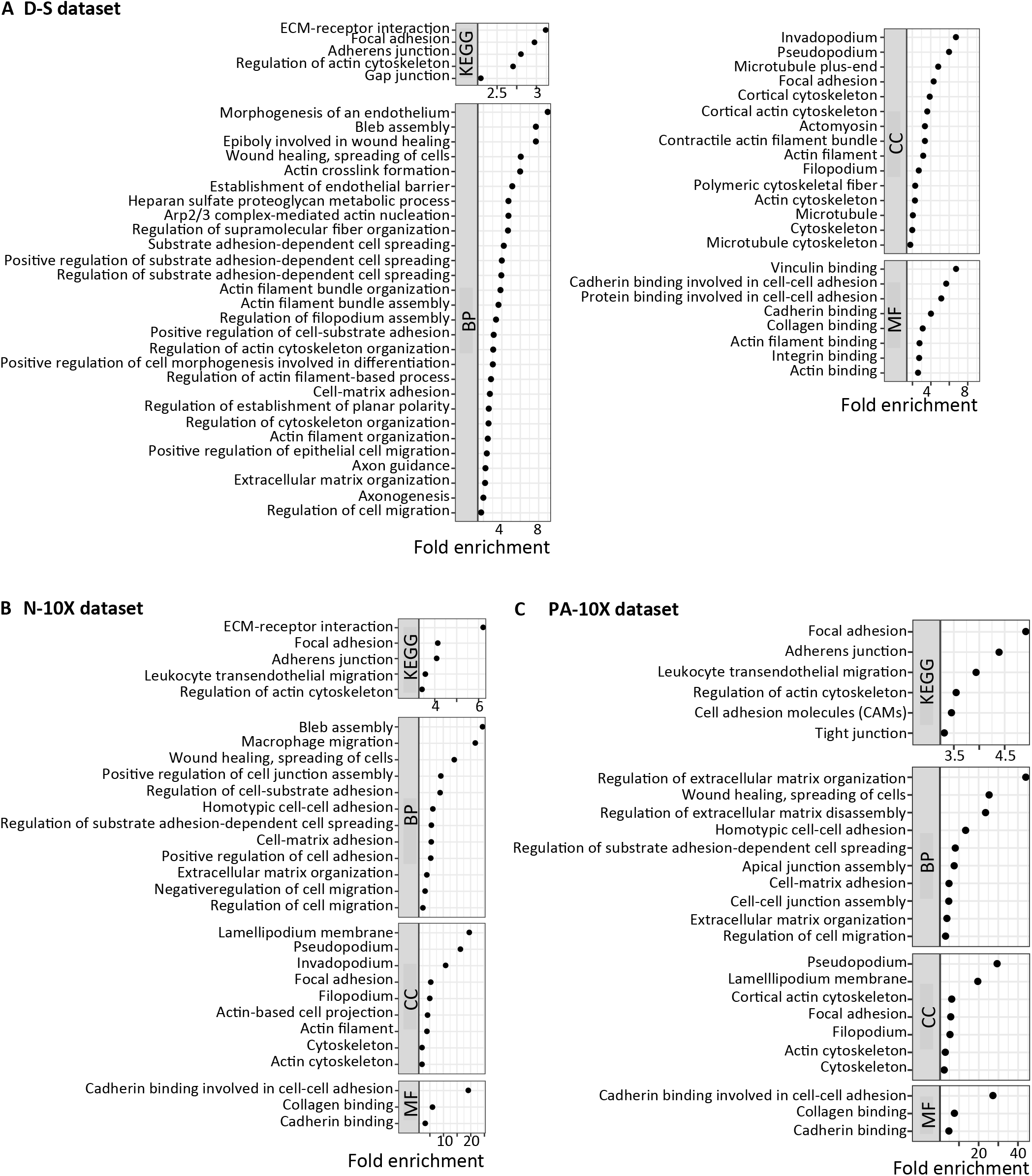
Motility-related terms highlighted by ontology analysis of genes overexpressed in M^HIGH^ versus M^LOW^ cells from independent datasets. Dot plot representation of enriched terms related to motility. Analyses performed using genes overexpressed with fold change >2 (A, D-S dataset, 1486 genes) and (B, N-10X dataset, 600 genes), and fold change >1.5 (C, PA-10X dataset, 338 genes). Benjamini-Hochberg (BH)-adjusted p-value < 0.05. KEGG: Kyoto Encyclopedia of Genes and Genomes, BP: Biological processes, CC: Cellular components, MF: Molecular functions.

**Supplementary Figure 3, related to Fig.1.**
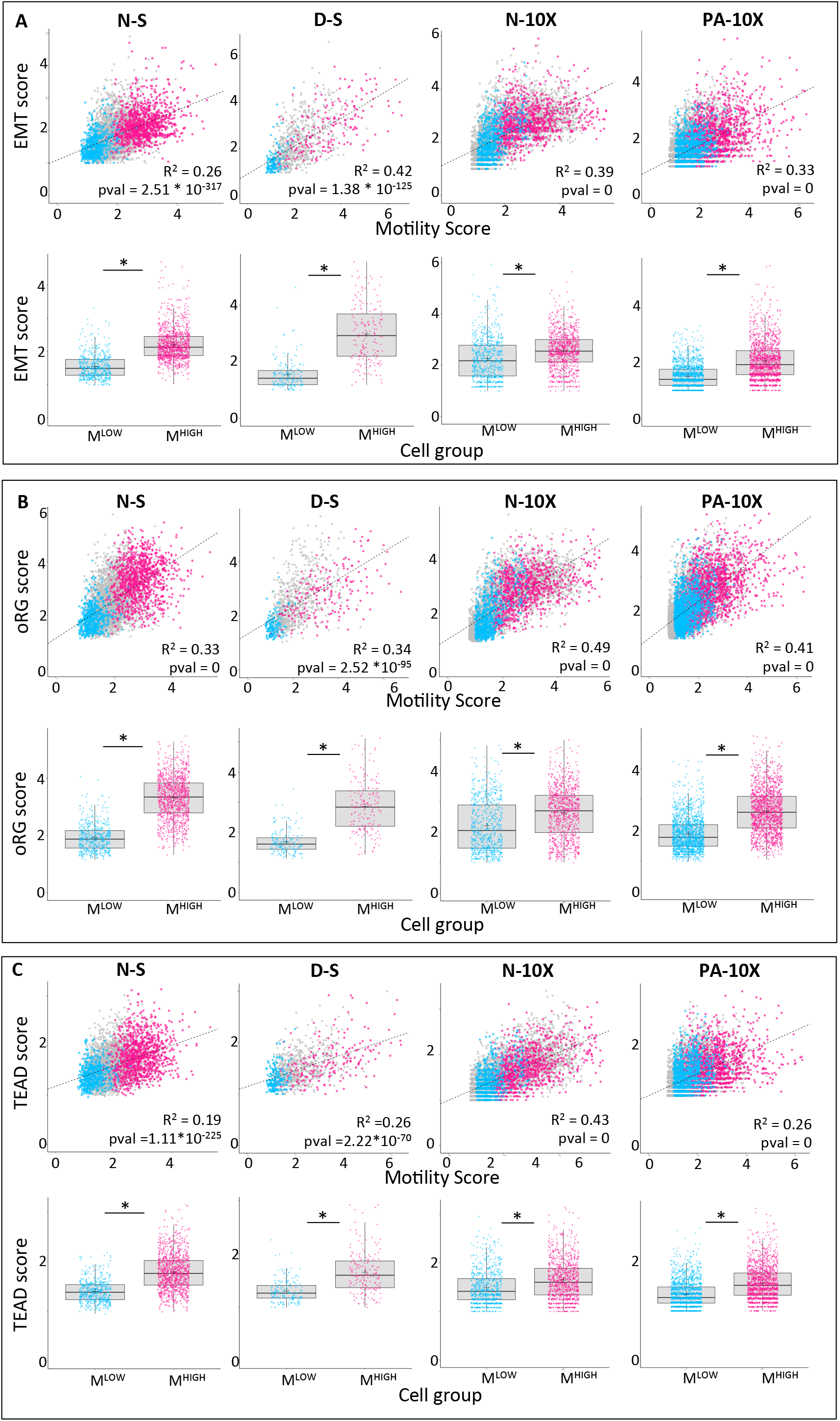
Enrichment in EMT (A), oRG (B) and TEAD (C) gene modules previously associated with glioblastoma cell motility in M^HIGH^ cells. EMT: genes associated with epithelio-mesenchymal transformation and GB cell motility. oRG: genes signing for outer Radial Glia (oRG)-like malignant cell population with increased invasive behavior in GB. TEAD: TEAD1-regulated genes involved in GB cell motility. Upper subpanels: Linear regression models between motility score and EMT, oRG and TEAD scores. p < 0.0001. Lower subpanels: Higher EMT, oRG and TEAD scores in M^HIGH^ versus M^LOW^ cells. *: p < 0.0001, Mann-Whitney test.

**Supplementary Figure 4, related to Fig.3-5.**
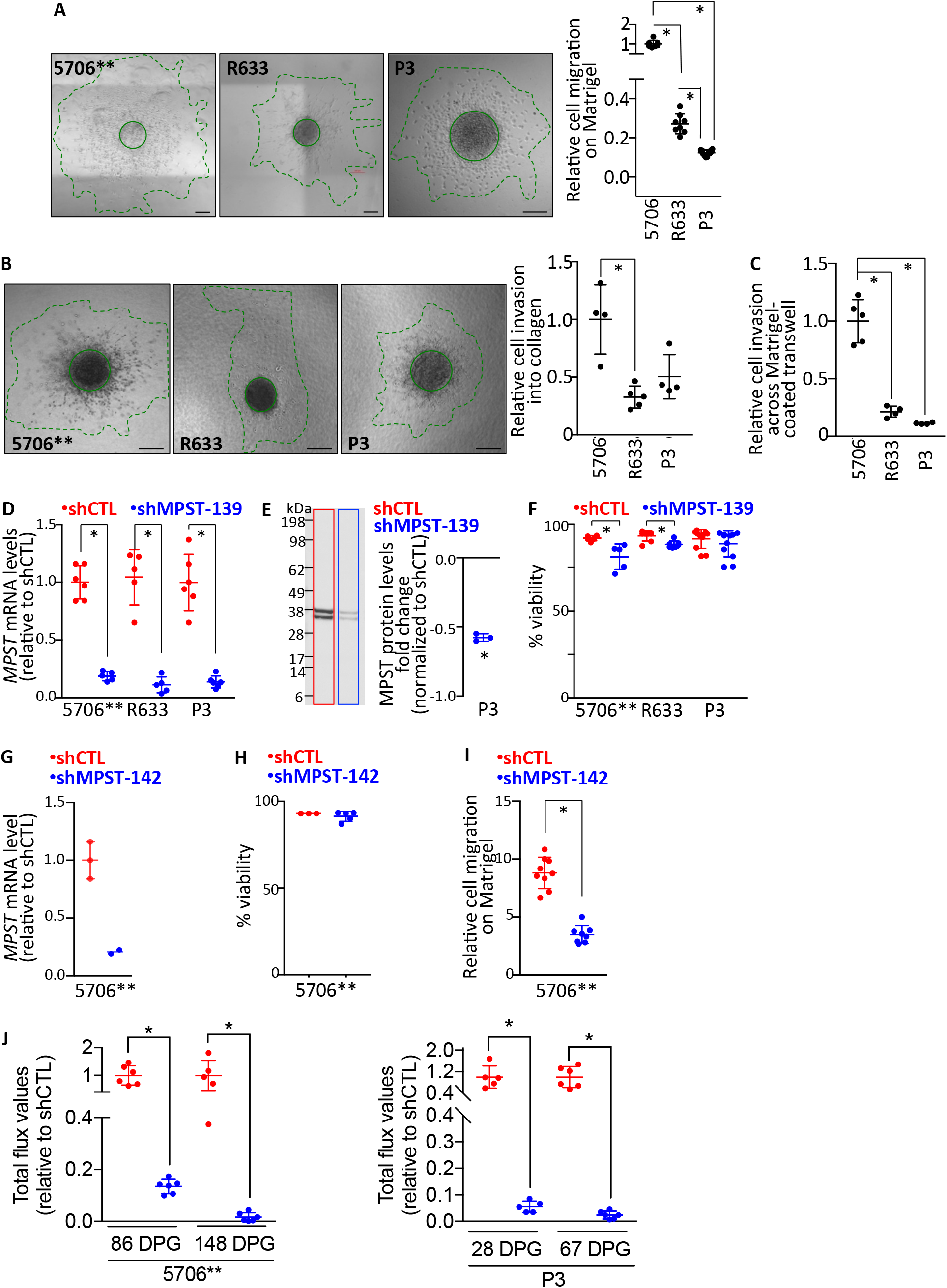
**A-C. Comparative migratory and invasive properties between glioblastoma PDC. A.** Cell migration on Matrigel assessed after 24h (5706**, R633 and P3). The microphotographs illustrate examples for each PDC. Scale bar = 200μm. A solid and dotted line delineate the spheroid core and the migration area, respectively. The dot plot depicts the quantification of the migration. Mean ± SD, n = 6-12 independent biological samples, *: p < 0.05, Tukey’s multiple comparisons test. **B.** Cell invasion into collagen assessed after 40h (5706**) and 23-25h (R633, P3). Microphotographs provide examples of a cell invasion assay for each PDC. Scale bars = 200μm. A solid and dotted line delineate the spheroid core and the invasion area, respectively. The bar graph depicts the quantification of the invasion. Mean ± SD, n = 4-5 independent biological samples, *: p < 0.05, Tukey’s multiple comparisons test. **C.** Quantification of cell invasion across Matrigel-coated transwells after 24 hours. 5706**, R633, and P3 PDC. Mean ± SD, n = 4-5 independent biological samples. **D-I. *MPST* knockdown. D.** Decreased *MPST* mRNA levels in PDC expressing *shMPST-139.* Mean ± SD, n = 5-6 independent biological samples. *: p < 0.05, Mann-Whitney test. **E.** Decreased MPST protein levels in *shMPST* expressing PDC. Western Blot analysis. MPST MW: 33/35 kDa. *: p < 0.05, one-sample t-test. **F.** No major impact of *MPST* knockdown on cell viability. Cell viability assessed using Trypan Blue exclusion test. Mean ± SD, n = 5-12 independent biological samples, *: p < 0.05, Mann-Whitney test. 5706**, R633 and P3 PDC. **G.** Decreased *MPST* mRNA levels in shMPST142-PDC compared to shControl-PDC. Q-PCR assay. Mean ± SD, n = 2-3 independent biological samples. **H.** No change in cell viability upon shMPST-142 expression assessed using Trypan Blue exclusion test. Mean ± SD, n = 3-5 independent biological samples, unpaired t-test with Welch correction. **I.** Decreased cell migration upon *MPST* knockdown. Cell migration assessed on Matrigel 7h post-seeding. Mean ± SD, n = 8-9 independent biological samples, *: p < 0.05, Mann-Whitney test. Quantification of cell invasion across Matrigel-coated transwells after 24 hours. 5706**, R633, and P3 PDC. Mean ± SD, n = 3-5 independent biological samples. **J. *MPST* knockdown decreases tumor burden until experiment end-points.** DPG: days post-graft. Mean ± SD, *: p < 0.01, Mann-Whitney test. 5706**-PDC: 86 DPG, n = 6 mice per group; 148 DPG, n = 5 shCTL and n = 6 shMPST. P3-PDC: 28 DPG, n = 5 mice per group; 67 DPG, n = 6 mice per group.

**Supplementary Figure 5, related to Methods.**
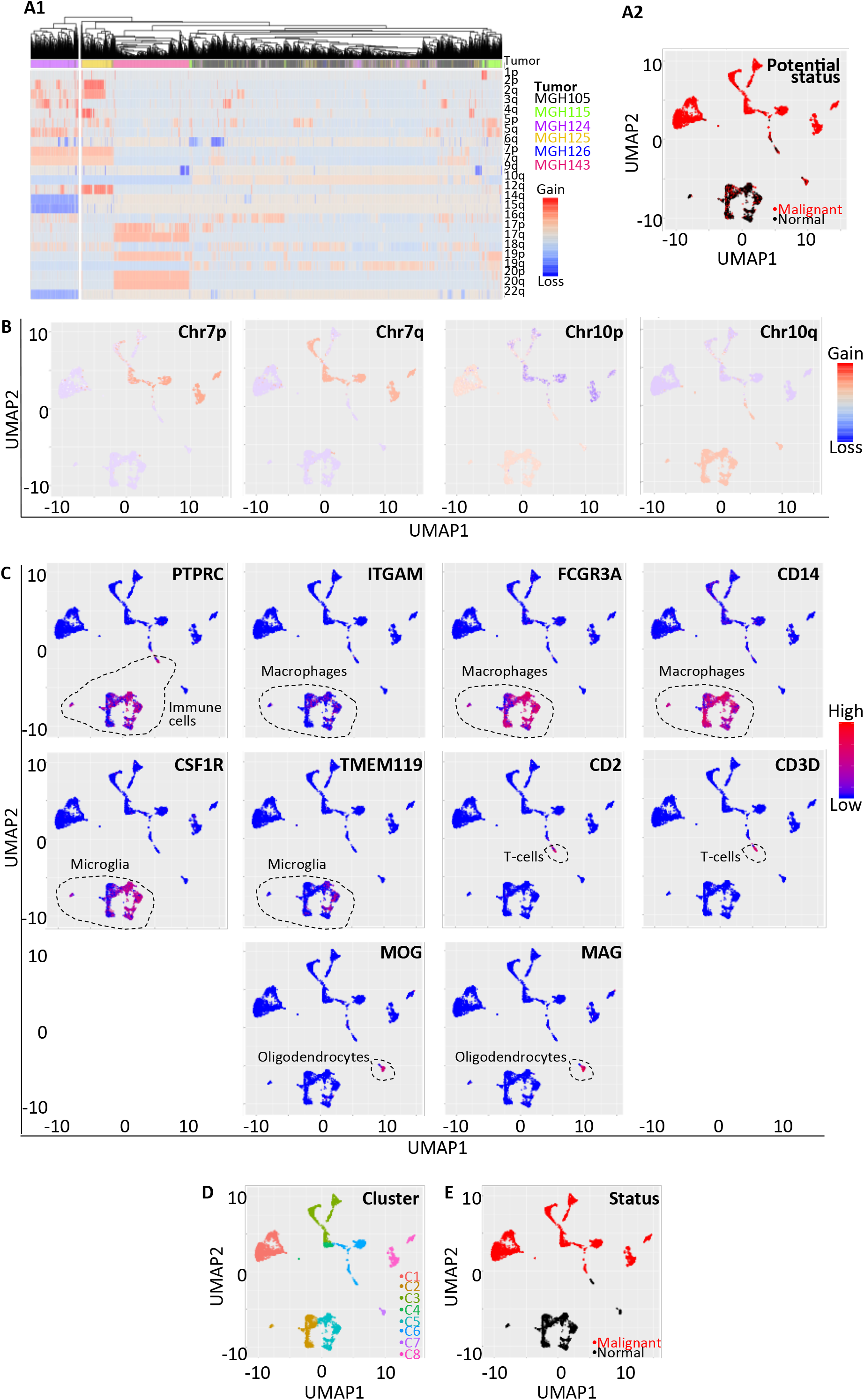
Identification of malignant and normal cells in N-10X dataset. **A.** Cell clustering based on GMM-based CNV predictions. Heatmap representation of CNV (copy number variations) predictions (A1). Potential malignancy status assigned, following cell clustering based on CNV predictions, highlighted on UMAP representation (A2). **B.** CNV predictions at canonical glioblastoma loci (Chr7 and 10). UMAP representation. **C.** Expression of marker genes of normal cell types. Pan-immune cells (PTPRC), macrophages (ITGAM, FCGR3A, CD14), microglia (CSF1R, TMEM119), T-cells (CD2, CD3D) and oligodendrocytes (MOG, MAG). UMAP representation. **D.** Cell clusters identified based on their repartition on UMAP plot. Hierarchical clustering followed by K-means clustering on UMAP components. **E.** Cell malignancy status assigned based on CNV status prediction, marker gene expression and clustering results. UMAP representation.

**Supplementary Figure 6, related to Methods.**
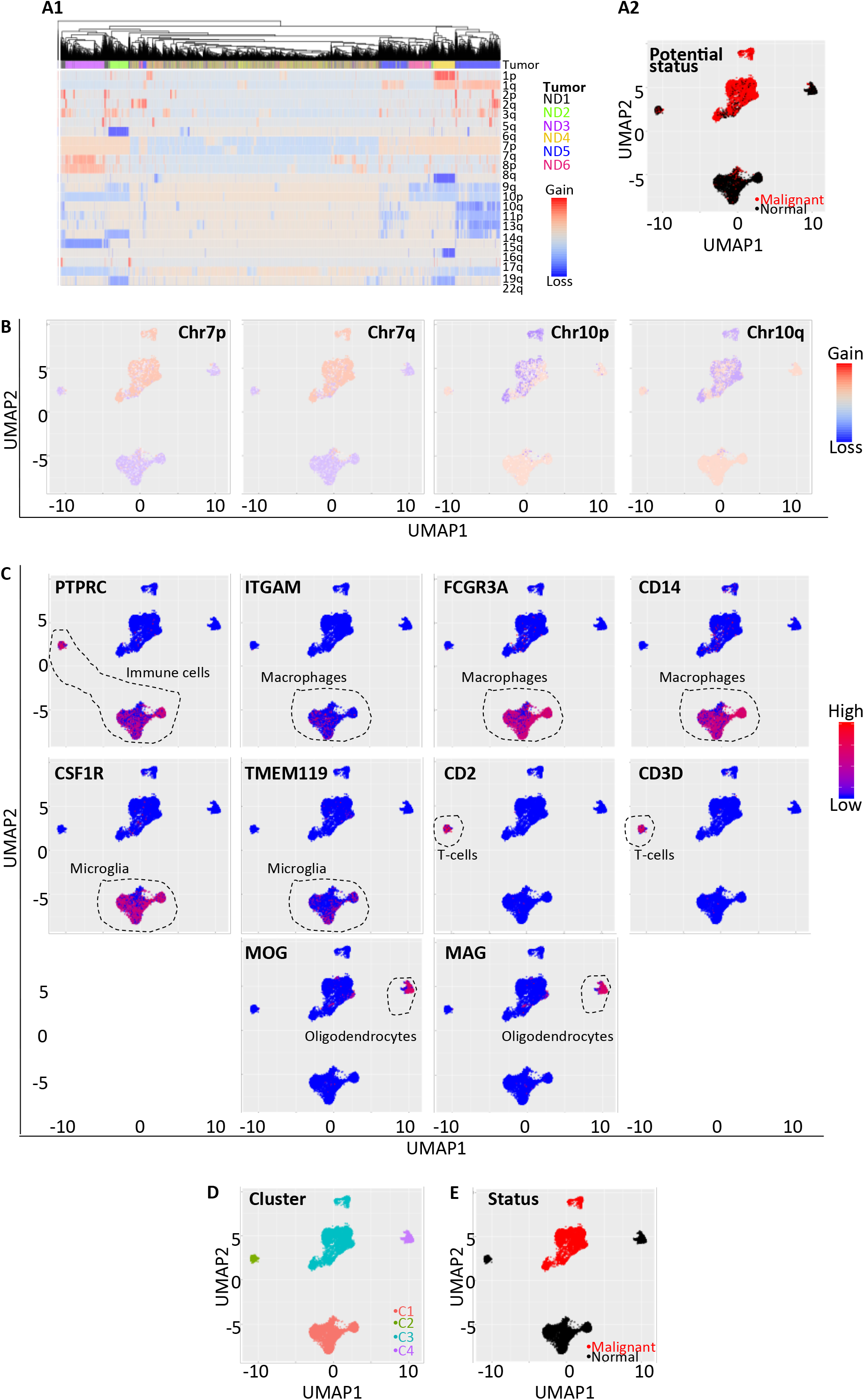
Identification of malignant and normal cells in PA-10X dataset. **A.** Cell clustering based on GMM-based CNV predictions. Heatmap representation of CNV (copy number variations) predictions (A1). Potential malignancy status assigned, following cell clustering based on CNV predictions, highlighted on UMAP representation (A2). **B.** CNV predictions at canonical glioblastoma loci (Chr7 and 10). UMAP representation. **C.** Expression of marker genes of normal cell types. Pan-immune cells (PTPRC), macrophages (ITGAM, FCGR3A, CD14), microglia (CSF1R, TMEM119), T-cells (CD2, CD3D) and oligodendrocytes (MOG, MAG). UMAP representation. **D.** Cell clusters identified based on their repartition on UMAP plot. Hierarchical clustering followed by K-means clustering on UMAP components. **E.** Cell malignancy status assigned based on CNV status prediction, marker gene expression and clustering results. UMAP representation.

**Supplementary Datafile 1**: Rscripts used in this study

## Supplementary Tables

**Supplementary Table 1, related to Fig 1 and Supplementary Fig. 1. Motility signature genes**

Sheet 1: description of the contents

Sheet2: list of genes in the motility signature and the articles experimentally demonstrating its implication in GB cell motility. Sheet3: results of the Pearson correlation analysis between the motility signature genes in the N-S dataset.

**Supplementary Table 2, related to Fig. 1 and Supplementary Fig. 1–3. Differential expression analysis results between M^HIGH^ and M^LOW^ cells (sheets 2-6) and list of genes in the oRG, EMT and TEAD-induced migration signatures (Sheet 7).**

Sheet 1: description of the contents

Sheet2: list of genes differentially expressed between M^HIGH^ and M^LOW^ cells from N-S dataset

Sheet3: list of genes differentially expressed between M^HIGH^ and M^LOW^ cells from D-S dataset

Sheet4: list of genes differentially expressed between M^HIGH^ and M^LOW^ cells from N-10X dataset

Sheet5: list of genes differentially expressed between M^HIGH^ and M^LOW^ cells from PA-10X dataset

Sheet6: list of genes overexpressed in M^HIGH^ versus M^LOW^ cells from the 4 datasets

Sheet7: list of genes in the oRG, EMT and TEAD-induced migration signatures used

**Supplementary Table 3, related to Fig. 1 and Supplementary Fig. 2. Results of gene ontology (GO) and KEGG pathway analyses preformed with the list of genes overexpressed in M^HIGH^ versus M^LOW^ cells**

Sheet 1: description of the contents

Sheet 2: results obtained using genes overexpressed in M^HIGH^ versus M^LOW^ cells with FC >2 (N-S dataset)

Sheet 3: results obtained using genes overexpressed in M^HIGH^ versus M^LOW^ cells with FC >2 (D-S dataset)

Sheet 4: results obtained using genes overexpressed in M^HIGH^ versus M^LOW^ cells with FC >2 (N-10X dataset)

Sheet 5: results obtained using genes overexpressed in M^HIGH^ versus M^LOW^ cells with FC >1.5 (PA-10X dataset)

**Supplementary Table 4, related to Fig. 2. KEGG pathway analysis of metabolism genes overexpressed in glioblastoma cells with high motile potential**

Sheet 1: description of the contents

“Sheet2: list of metabolism genes upregulated in M-HIGH versus M-LOW cells from each dataset”

Sheet3: results of the KEGG pathway analysis performed on metabolism genes upregulated in M-HIGH versus M-LOW cells from N-S dataset (BH-adj pval < 0.05)

Sheet4: results of the KEGG pathway analysis performed on metabolism genes upregulated in M-HIGH versus M-LOW cells from D-S dataset (BH-adj pval < 0.05)

Sheet5: results of the KEGG pathway analysis performed on metabolism genes upregulated in M-HIGH versus M-LOW cells from N-10X dataset (BH-adj pval < 0.05)

Sheet6: results of the KEGG pathway analysis performed on metabolism genes upregulated in M-HIGH versus M-LOW cells from PA-10X dataset (BH-adj pval < 0.05)

**Supplementary Table 5, related to Fig. 2. Metabolism genes overexpressed in S3-S1 branch compared to each other branch**

**Supplementary Table 6, related to methods**

## References

Abdollahi Govar A, Toro G, Szaniszlo P, Pavlidou A, Bibli SI, et al. 2020. 3-Mercaptopyruvate sulfurtransferase supports endothelial cell angiogenesis and bioenergetics. Br J Pharmacol 177: 866–83

Armento A, Ehlers J, Schötterl S, Naumann U. 2017. Molecular Mechanisms of Glioma Cell Motility. Glioblastoma [Internet]

Augsburger F, Randi EB, Jendly M, Ascencao K, Dilek N, Szabo C. 2020. Role of 3-Mercaptopyruvate Sulfurtransferase in the Regulation of Proliferation, Migration, and Bioenergetics in Murine Colon Cancer Cells. Biomolecules 10

Bhaduri A, Di Lullo E, Jung D, Muller S, Crouch EE, et al. 2020. Outer Radial Glia-like Cancer Stem Cells Contribute to Heterogeneity of Glioblastoma. Cell Stem Cell 26: 48–63 e6

Caino MC, Ghosh JC, Chae YC, Vaira V, Rivadeneira DB, et al. 2015. PI3K therapy reprograms mitochondrial trafficking to fuel tumor cell invasion. Proc Natl Acad Sci U S A 112: 8638–43

Carro MS, Lim WK, Alvarez MJ, Bollo RJ, Zhao X, et al. 2010. The transcriptional network for mesenchymal transformation of brain tumours. Nature 463: 318–25

Chen EY, Tan CM, Kou Y, Duan Q, Wang Z, et al. 2013. Enrichr: interactive and collaborative HTML5 gene list enrichment analysis tool. BMC Bioinformatics 14: 128

Chen H, Albergante L, Hsu JY, Lareau CA, Lo Bosco G, et al. 2019. Single-cell trajectories reconstruction, exploration and mapping of omics data with STREAM. Nat Commun 10: 1903

Cooper AJ, Shurubor YI, Dorai T, Pinto JT, Isakova EP, et al. 2016. omega-Amidase: an underappreciated, but important enzyme in L-glutamine and L-asparagine metabolism; relevance to sulfur and nitrogen metabolism, tumor biology and hyperammonemic diseases. Amino Acids 48: 1–20

Courte J, Renault R, Jan A, Viovy JL, Peyrin JM, Villard C. 2018. Reconstruction of directed neuronal networks in a microfluidic device with asymmetric microchannels. Methods Cell Biol 148: 71–95

Couturier CP, Ayyadhury S, Le PU, Nadaf J, Monlong J, et al. 2020. Single-cell RNA-seq reveals that glioblastoma recapitulates a normal neurodevelopmental hierarchy. Nat Commun 11: 3406

Cuddapah VA, Robel S, Watkins S, Sontheimer H. 2014. A neurocentric perspective on glioma invasion. Nat Rev Neurosci 15: 45565

Darmanis S, Sloan SA, Croote D, Mignardi M, Chernikova S, et al. 2017. Single-Cell RNA-Seq Analysis of Infiltrating Neoplastic Cells at the Migrating Front of Human Glioblastoma. Cell Rep 21: 1399–410

Daubon T, Leon C, Clarke K, Andrique L, Salabert L, et al. 2019. Deciphering the complex role of thrombospondin-1 in glioblastoma development. Nat Commun 10: 1146

de Gooijer MC, Guillen Navarro M, Bernards R, Wurdinger T, van Tellingen O. 2018. An Experimenter’s Guide to Glioblastoma Invasion Pathways. Trends Mol Med 24: 763–80

Dustin CM, Heppner DE, Lin MJ, van der Vliet A. 2020. Redox regulation of tyrosine kinase signalling: more than meets the eye. J Biochem 167: 151–63

El-Habr EA, Dubois LG, Burel-Vandenbos F, Bogeas A, Lipecka J, et al. 2017. A driver role for GABA metabolism in controlling stem and proliferative cell state through GHB production in glioma. Acta Neuropathol 133: 645–60

Eskilsson E, Rosland GV, Talasila KM, Knappskog S, Keunen O, et al. 2016. EGFRvIII mutations can emerge as late and heterogenous events in glioblastoma development and promote angiogenesis through Src activation. Neuro Oncol 18: 1644–55

Filipovic MR, Zivanovic J, Alvarez B, Banerjee R. 2018. Chemical Biology of H2S Signaling through Persulfidation. Chem Rev 118: 1253–337

Flavahan WA, Wu Q, Hitomi M, Rahim N, Kim Y, et al. 2013. Brain tumor initiating cells adapt to restricted nutrition through preferential glucose uptake. Nat Neurosci 16: 1373–82

Frolov A, Evans IM, Li N, Sidlauskas K, Paliashvili K, et al. 2016. Imatinib and Nilotinib increase glioblastoma cell invasion via Abl-independent stimulation of p130Cas and FAK signalling. Sci Rep 6: 27378

Gagliardi F, Narayanan A, Mortini P. 2017. SPARCL1 a novel player in cancer biology. Crit Rev Oncol Hematol 109: 63–68

Gao XH, Krokowski D, Guan BJ, Bederman I, Majumder M, et al. 2015. Quantitative H2S-mediated protein sulfhydration reveals metabolic reprogramming during the integrated stress response. Elife 4: e10067

Garcia JH, Jain S, Aghi MK. 2021. Metabolic Drivers of Invasion in Glioblastoma. Front Cell Dev Biol 9: 683276

Gu Z, Eils R, Schlesner M. 2016. Complex heatmaps reveal patterns and correlations in multidimensional genomic data. Bioinformatics 32: 2847–9

Gu Z, Gu L, Eils R, Schlesner M, Brors B. 2014. circlize Implements and enhances circular visualization in R. Bioinformatics 30: 2811–2

Guyon J, Andrique L, Pujol N, Rosland GV, Recher G, et al. 2020. A 3D Spheroid Model for Glioblastoma. J Vis Exp

Hambardzumyan D, Bergers G. 2015. Glioblastoma: Defining Tumor Niches. Trends Cancer 1: 252–65

Hanaoka K, Sasakura K, Suwanai Y, Toma-Fukai S, Shimamoto K, et al. 2017. Discovery and Mechanistic Characterization of Selective Inhibitors of H2S-producing Enzyme: 3-Mercaptopyruvate Sulfurtransferase (3MST) Targeting Active-site Cysteine Persulfide. Sci Rep 7: 40227

Huber SM, Butz L, Stegen B, Klumpp D, Braun N, et al. 2013. Ionizing radiation, ion transports, and radioresistance of cancer cells. Front Physiol 4: 212

Husson F, Josse J, J. P. 2010. Principal component methods – hierarchical clustering – partitional clustering: why would we need to choose for visualizing data?.

Ida T, Sawa T, Ihara H, Tsuchiya Y, Watanabe Y, et al. 2014. Reactive cysteine persulfides and S-polythiolation regulate oxidative stress and redox signaling. Proc Natl Acad Sci U S A 111: 7606–11

Jacob F, Salinas RD, Zhang DY, Nguyen PTT, Schnoll JG, et al. 2020. A Patient-Derived Glioblastoma Organoid Model and Biobank Recapitulates Inter- and Intra-tumoral Heterogeneity. Cell 180: 188–204 e22

Ji J, Xu R, Ding K, Bao G, Zhang X, et al. 2019. Long Noncoding RNA SChLAP1 Forms a Growth-Promoting Complex with HNRNPL in Human Glioblastoma through Stabilization of ACTN4 and Activation of NF-kappaB Signaling. Clin Cancer Res 25: 6868–81

Joseph JV, Conroy S, Tomar T, Eggens-Meijer E, Bhat K, et al. 2014. TGF-beta is an inducer of ZEB1-dependent mesenchymal transdifferentiation in glioblastoma that is associated with tumor invasion. Cell Death Dis 5: e1443

Joseph JV, Magaut CR, Storevik S, Geraldo LH, Mathivet T, et al. 2021. TGF-beta promotes microtube formation in glioblastoma through thrombospondin 1. Neuro Oncol

Kanehisa M, Furumichi M, Tanabe M, Sato Y, Morishima K. 2017. KEGG: new perspectives on genomes, pathways, diseases and drugs. Nucleic Acids Res 45: D353–D61

Kang W, Kim SH, Cho HJ, Jin J, Lee J, et al. 2015. Talin1 targeting potentiates anti-angiogenic therapy by attenuating invasion and stem-like features of glioblastoma multiforme. Oncotarget 6: 27239–51

Kimura Y, Koike S, Shibuya N, Lefer D, Ogasawara Y, Kimura H. 2017. 3-Mercaptopyruvate sulfurtransferase produces potential redox regulators cysteine- and glutathione-persulfide (Cys-SSH and GSSH) together with signaling molecules H2S2, H2S3 and H2S. Sci Rep 7: 10459

Kuleshov MV, Jones MR, Rouillard AD, Fernandez NF, Duan Q, et al. 2016. Enrichr: a comprehensive gene set enrichment analysis web server 2016 update. Nucleic Acids Res 44: W90–7

Lee JK, Wang J, Sa JK, Ladewig E, Lee HO, et al. 2017. Spatiotemporal genomic architecture informs precision oncology in glioblastoma. Nat Genet 49: 594–99

Ljubimova JY, Fujita M, Khazenzon NM, Ljubimov AV, Black KL. 2006. Changes in laminin isoforms associated with brain tumor invasion and angiogenesis. Front Biosci 11: 81–8

Lopez-Colome AM, Lee-Rivera I, Benavides-Hidalgo R, Lopez E. 2017. Paxillin: a crossroad in pathological cell migration. J Hematol Oncol 10: 50

Lu KV, Chang JP, Parachoniak CA, Pandika MM, Aghi MK, et al. 2012. VEGF inhibits tumor cell invasion and mesenchymal transition through a MET/VEGFR2 complex. Cancer Cell 22: 21–35

Manning C, Raghavan P, Schütze H. 2008. Flat clustering. Cambridge University Press: 321–45

Montana V, Sontheimer H. 2011. Bradykinin promotes the chemotactic invasion of primary brain tumors. J Neurosci 31: 4858–67

Muller S, Cho A, Liu SJ, Lim DA, Diaz A. 2018. CONICS integrates scRNA-seq with DNA sequencing to map gene expression to tumor sub-clones. Bioinformatics 34: 3217–19

Murphy B, Bhattacharya R, Mukherjee P. 2019. Hydrogen sulfide signaling in mitochondria and disease. FASEB J 33: 13098–125

Mustafa AK, Gadalla MM, Sen N, Kim S, Mu W, et al. 2009. H2S signals through protein S-sulfhydration. Sci Signal 2: ra72

Neftel C, Laffy J, Filbin MG, Hara T, Shore ME, et al. 2019. An Integrative Model of Cellular States, Plasticity, and Genetics for Glioblastoma. Cell 178: 835–49 e21

Oizel K, Chauvin C, Oliver L, Gratas C, Geraldo F, et al. 2017. Efficient Mitochondrial Glutamine Targeting Prevails Over Glioblastoma Metabolic Plasticity. Clin Cancer Res 23: 6292–304

Osswald M, Jung E, Sahm F, Solecki G, Venkataramani V, et al. 2015. Brain tumour cells interconnect to a functional and resistant network. Nature 528: 93–8

Pedersen PH, Edvardsen K, Garcia-Cabrera I, Mahesparan R, Thorsen J, et al. 1995. Migratory patterns of lac-z transfected human glioma cells in the rat brain. Int J Cancer 62: 767–71

Pedre B, Dick TP. 2021. 3-Mercaptopyruvate sulfurtransferase: an enzyme at the crossroads of sulfane sulfur trafficking. Biol Chem 402: 223–37

Pombo Antunes AR, Scheyltjens I, Lodi F, Messiaen J, Antoranz A, et al. 2021. Single-cell profiling of myeloid cells in glioblastoma across species and disease stage reveals macrophage competition and specialization. Nat Neurosci 24: 595–610

Porporato PE, Payen VL, Perez-Escuredo J, De Saedeleer CJ, Danhier P, et al. 2014. A mitochondrial switch promotes tumor metastasis. Cell Rep 8: 754–66

Puchalski RB, Shah N, Miller J, Dalley R, Nomura SR, et al. 2018. An anatomic transcriptional atlas of human glioblastoma. Science 360: 660–63

Pudelek M, Krol K, Catapano J, Wrobel T, Czyz J, Ryszawy D. 2020. Epidermal Growth Factor (EGF) Augments the Invasive Potential of Human Glioblastoma Multiforme Cells via the Activation of Collaborative EGFR/ROS-Dependent Signaling. Int J Mol Sci 21

Renault-Mihara F, Beuvon F, Iturrioz X, Canton B, De Bouard S, et al. 2006. Phosphoprotein enriched in astrocytes-15 kDa expression inhibits astrocyte migration by a protein kinase C delta-dependent mechanism. Mol Biol Cell 17: 5141–52

Richards LM, Whitley OKN, MacLeod G, Cavalli FMG, Coutinho FJ, et al. 2021. Gradient of Developmental and Injury Response transcriptional states defines functional vulnerabilities underpinning glioblastoma heterogeneity. Nature Cancer 2: 157–73

Rosenberg S, Verreault M, Schmitt C, Guegan J, Guehennec J, et al. 2017. Multi-omics analysis of primary glioblastoma cell lines shows recapitulation of pivotal molecular features of parental tumors. Neuro Oncol 19: 219–28

Saurty-Seerunghen MS, Bellenger L, El-Habr EA, Delaunay V, Garnier D, et al. 2019. Capture at the single cell level of metabolic modules distinguishing aggressive and indolent glioblastoma cells. Acta Neuropathol Commun 7: 155

Scherer HJ. 1938. Structural development in gliomas. American Journal of Cancer 34: 333–51

Sen S, Dong M, Kumar S. 2009. Isoform-specific contributions of alpha-actinin to glioma cell mechanobiology. PLoS One 4: e8427

Silver DJ, Roversi GA, Bithi N, Wang SZ, Troike KM, et al. 2021. Severe consequences of a high-lipid diet include hydrogen sulfide dysfunction and enhanced aggression in glioblastoma. J Clin Invest

Snuderl M, Fazlollahi L, Le LP, Nitta M, Zhelyazkova BH, et al. 2011. Mosaic amplification of multiple receptor tyrosine kinase genes in glioblastoma. Cancer Cell 20: 810–7

Soneson C, Robinson MD. 2018. Bias, robustness and scalability in single-cell differential expression analysis. Nat Methods 15: 255–61

Sottoriva A, Spiteri I, Piccirillo SG, Touloumis A, Collins VP, et al. 2013. Intratumor heterogeneity in human glioblastoma reflects cancer evolutionary dynamics. Proc Natl Acad Sci U S A 110: 4009–14

Stupp R, Taillibert S, Kanner A, Read W, Steinberg D, et al. 2017. Effect of Tumor-Treating Fields Plus Maintenance Temozolomide vs Maintenance Temozolomide Alone on Survival in Patients With Glioblastoma: A Randomized Clinical Trial. JAMA 318: 2306–16

Sun LH, Yang FQ, Zhang CB, Wu YP, Liang JS, et al. 2017. Overexpression of Paxillin Correlates with Tumor Progression and Predicts Poor Survival in Glioblastoma. CNS Neurosci Ther 23: 69–75

Takano N, Sarfraz Y, Gilkes DM, Chaturvedi P, Xiang L, et al. 2014. Decreased expression of cystathionine beta-synthase promotes glioma tumorigenesis. Mol Cancer Res 12: 1398–406

Tao B, Wang R, Sun C, Zhu Y. 2017. 3-Mercaptopyruvate Sulfurtransferase, Not Cystathionine beta-Synthase Nor Cystathionine gamma-Lyase, Mediates Hypoxia-Induced Migration of Vascular Endothelial Cells. Front Pharmacol 8: 657

Tentler D, Lomert E, Novitskaya K, Barlev NA. 2019. Role of ACTN4 in Tumorigenesis, Metastasis, and EMT. Cells 8

Tome-Garcia J, Erfani P, Nudelman G, Tsankov AM, Katsyv I, et al. 2018. Analysis of chromatin accessibility uncovers TEAD1 as a regulator of migration in human glioblastoma. Nat Commun 9: 4020

Untereiner AA, Olah G, Modis K, Hellmich MR, Szabo C. 2017. H2S-induced S-sulfhydration of lactate dehydrogenase a (LDHA) stimulates cellular bioenergetics in HCT116 colon cancer cells. Biochem Pharmacol 136: 86–98

Vollmann-Zwerenz A, Leidgens V, Feliciello G, Klein CA, Hau P. 2020. Tumor Cell Invasion in Glioblastoma. Int J Mol Sci 21

Volovetz J, Berezovsky AD, Alban T, Chen Y, Lauko A, et al. 2020. Identifying conserved molecular targets required for cell migration of glioblastoma cancer stem cells. Cell Death Dis 11: 152

Wang R, Sharma R, Shen X, Laughney AM, Funato K, et al. 2020. Adult Human Glioblastomas Harbor Radial Glia-like Cells. Stem Cell Reports 14: 338–50

Watkins S, Robel S, Kimbrough IF, Robert SM, Ellis-Davies G, Sontheimer H. 2014. Disruption of astrocyte-vascular coupling and the blood-brain barrier by invading glioma cells. Nat Commun 5: 4196

Weng MS, Chang JH, Hung WY, Yang YC, Chien MH. 2018. The interplay of reactive oxygen species and the epidermal growth factor receptor in tumor progression and drug resistance. J Exp Clin Cancer Res 37: 61

Xia S, Lal B, Tung B, Wang S, Goodwin CR, Laterra J. 2016. Tumor microenvironment tenascin-C promotes glioblastoma invasion and negatively regulates tumor proliferation. Neuro Oncol 18: 507–17

Xie Z, Bailey A, Kuleshov MV, Clarke DJB, Evangelista JE, et al. 2021. Gene Set Knowledge Discovery with Enrichr. Curr Protoc 1: e90

Zivanovic J, Kouroussis E, Kohl JB, Adhikari B, Bursac B, et al. 2019. Selective Persulfide Detection Reveals Evolutionarily Conserved Antiaging Effects of S-Sulfhydration. Cell Metab 30: 1152–70 e13

Zuhra K, Tome CS, Forte E, Vicente JB, Giuffre A. 2021. The multifaceted roles of sulfane sulfur species in cancer-associated processes. Biochim Biophys Acta Bioenerg 1862: 148338

